# Concussive Injury in Human Forebrain Organoids Alters Neurogenesis and Neuroimmune Signaling

**DOI:** 10.64898/2026.10.29.762930

**Authors:** Razieh Jaberi, Prince D Okoro, Christopher Taitano-Johnson, Erick Contreras, Rhea Chidambaram, Piunik Oganian Babakhanins, Iman Noshadi, Nicole I zur Nieden, Vijayalakshmi Santhakumar

## Abstract

Traumatic brain injury (TBI) is a major cause of long-term neurological disability, yet the mechanisms underlying human-specific injury responses remain poorly understood. Rodent models of TBI have yielded significant insights into mechanisms and pathophysiology of TBI but may not capture human-specific developmental and molecular responses. Here, we establish a human forebrain organoid (hFBO) injury paradigm to examine developmental stage and region-specific responses to concussive injury. Dorsal and ventral hFBOs, enriched in glutamatergic and GABAergic neurons, respectively, showed progressive physiological maturation with rudimentary firing characteristics at 80 days in vitro (DIV) and spontaneous synaptic activity by 150 DIV. Injury to dorsal and ventral hFBOs at 25 DIV, when neural progenitors dominate, caused cell death that scaled with severity and increased expression of neural proliferation markers and TLR4/NF-κB associated neuroimmune signaling. Injury to physiologically mature organoids at 150 DIV produced severity-dependent acute intracellular calcium dysregulation. Together, these findings establish a human organoid-based platform that captures developmental stage-dependent neurogenic, molecular, and cellular dysfunction in responses to concussive injury.

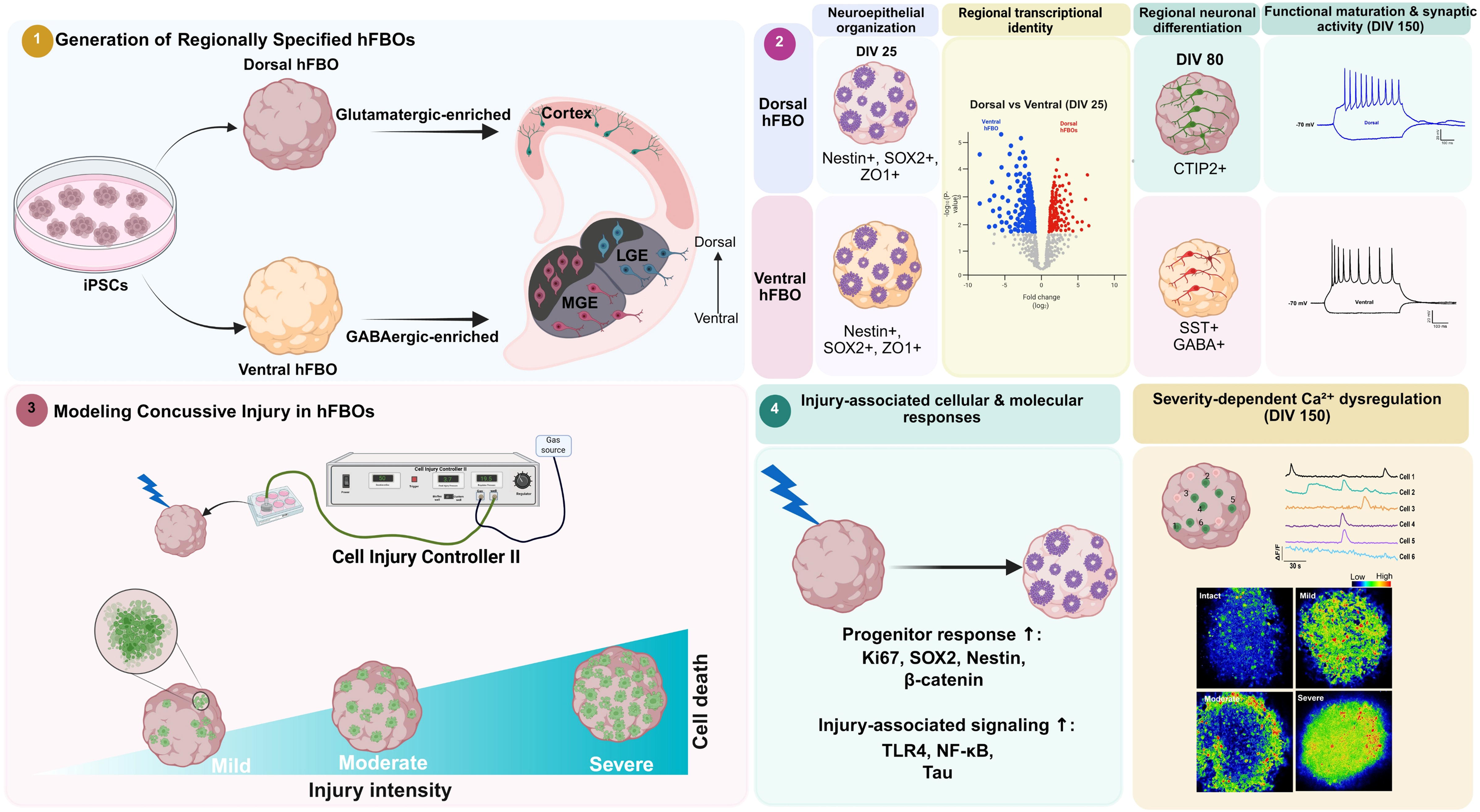

**Highlights:**

- Injury severity drives cell death in human forebrain organoids (hFBO)
- Developmental stage, but not regional identity, regulates injury-induced cell death
- Early injury increases neural progenitor proliferation and neuroimmune signaling
- Dorsal and ventral hFBOs show regional cellular specification by 80 DIV and active mature circuits by 150 DIV
- Injury at 150 DIV induces severity-dependent calcium dysregulation

## Introduction

Traumatic brain injury (TBI) is a major public health issue, affecting millions of individuals annually and frequently resulting in long-term cognitive and emotional impairments (Maas et al. 2022; Haarbauer-Krupa et al. 2021). Despite substantial advances in TBI research using rodent models, translation of mechanistic findings into effective human therapies remains a formidable challenge (Xiong, Mahmood, and Chopp 2013; Zhao et al. 2023). Rodent studies have provided critical insights into the roles of neuroinflammation, cell death, and circuit dysfunction in TBI pathology (Hamilton and Santhakumar 2017; Hunt, Boychuk, and Smith 2013; Pitkänen et al. 2009; Neuberger et al. 2019); however, these models fail to fully capture the molecular and cellular complexity of human concussive brain trauma (Swingler et al. 2023; S. H. Kim and Chang 2023; Lancaster and Knoblich 2014). Fundamental interspecies differences in development, cortical organization, gene expression, and regenerative responses highlight the need for human-relevant experimental systems to bridge this translational gap. Although animal-based and human-derived two- and three-dimensional *in vitro* models have advanced our understanding of TBI mechanisms, key features of highly prevalent concussive brain injury remain insufficiently modeled in human brain organoids (Smith and Meaney 2000; Redell et al. 2020; Ngwenya and Danzer 2019; Bielefeld et al. 2024). Concussive injury alters neurogenesis by modulating progenitor survival, proliferation, and differentiation (Clark et al. 2021; Neuberger, Swietek, et al. 2017); however, the molecular pathways driving these changes and their relevance to human tissue remain unclear (Clark et al. 2021). In more mature circuits, concussive injury also induces persistent alterations in neuronal excitability and microcircuit function that contribute to adverse long-term neurological outcomes (Bolkvadze and Pitkänen 2012; Korgaonkar, Li, et al. 2020; Neuberger et al. 2019). A versatile human-tissue model that captures developmental processes from neurogenesis to maturation of cellular physiology and circuit function is essential for understanding a broad spectrum of injury mechanisms in the developing brain. Concussive injuries are associated with a multitude of cellular and circuit changes, including alterations in neurogenesis and network excitability early after injury (Sun 2014; Witkowski et al. 2019). The early changes after trauma are driven by changes in cell signaling pathways, inflammatory responses, and microcircuit plasticity and shape both recovery and long-term neuropathology (Santhakumar et al. 2000; Arizanovska et al. 2025; Bonislawski, Schwarzbach, and Cohen 2007; Kang et al. 2022; Korgaonkar, Nguyen, et al. 2020; Neuberger, Gupta, et al. 2017). Furthermore, conventional neural culture systems lacking defined regional identity may not adequately capture region-specific cellular responses to traumatic injury. In contrast, regionally specified human forebrain organoids recapitulate distinct dorsal and ventral developmental programs, providing an opportunity to investigate injury responses across glutamatergic- and GABAergic-enriched neural populations (Cederquist et al. 2019; Birey et al. 2017). The three-dimensional, self-organized organoid systems provide a powerful human-relevant complement to *in vivo* animal models. The 3D human brain organoids recapitulate unique aspects of human brain development (Sloan, Andersen, Pa, et al. 2018; Eichmüller and Knoblich 2022; Adlakha 2023), can be differentiated into regionally specific structures (Xiang, Cakir, and Park 2025) and can serve as a vital bridge between preclinical and translational TBI research (Hamilton and Santhakumar 2017; Lai et al. 2024; Sirtori et al. 2025). To address the limitations of existing human tissue TBI models and enable mechanistic studies of human-specific responses to concussive brain injury, we developed a pressure-based concussive injury paradigm in hFBOs using the Cell Injury Controller II (CICII), typically used to model 2D stretch injury (Ellis et al. 1995). While stretch-, shear-, and ultrasound-based 2D and 3D human *in vitro* tissue injury models have provided important insights into injury-induced neuronal damage, most lack the three-dimensional cytoarchitecture, developmental patterning and cellular diversity required to model the biomechanical, molecular, and circuit-level consequences of concussive trauma (Liaudanskaya et al. 2021; Pilipovi 2023; Geddes-klein, Schiffman, and Meaney 2006; Bano, Jeevanandam, and Tsenov 2025; Shaughness 2021). Human brain organoids offer a self-organized, three-dimensional system that recapitulates key aspects of early human cortical development, regional patterning, and gene expression, enabling investigation of injury responses in a spatially relevant context (Li et al. 2025; Tamada and Muguruma 2024). By applying a brief, controlled pressure pulse to 3D organoid cultures, our model precisely modulates injury severity and captures features of concussive trauma that are difficult to reproduce in 2D systems. This approach allows direct comparison of injury responses across defined severities and provides a platform to examine how mechanical injury alters neuronal proliferation, survival, excitability, and circuit function in regionally specified human tissue (Ramirez et al. 2021; Lai et al. 2024). Importantly, this organoid-based *in vitro* concussive injury model can be studied in parallel with established rodent and porcine *in vivo* fluid percussion injury, offering a complementary framework to bridge mechanistic discoveries between animal models and human systems.

## Materials and Methods

### Human iPSC Culture and Maintenance

Two healthy control human induced pluripotent stem cell (iPSC) lines from healthy male and female donors (SCTi003-A, female, and SCTi004-A, male; STEMCELL Technologies) were used to generate organoids. Both iPSC lines were cultured under feeder-free conditions according to the manufacturer’s recommended protocols. Cells were maintained on tissue culture–treated plates pre-coated with human CellAdhere™ Vitronectin (0.5 µg/mL; purified from human serum; STEMCELL Technologies). SCTi003-A cells were cultured and maintained using mTeSR™ Plus medium, which was also used for thawing and routine culture. Cells were passaged enzymatically using ReLeSR™ (STEMCELL Technologies) at a 1:3 split ratio every 4–5 days upon reaching approximately 75% confluency (Sparks et al. 2022; Simorgh et al. 2023). All cultures were maintained at 37 °C in a humidified incubator with 5% CO₂. Cells were routinely tested and confirmed to be free of mycoplasma contamination. All experiments were performed using iPSCs at passage numbers below 10.

### Generation of Human Forebrain Organoids

Organoid differentiation was performed following established protocols based on dual-SMAD inhibition, with stage-specific small molecules and growth factors applied as outlined in the schematic (Figure 1A) (Sloan, Andersen, Pa, et al. 2018). Region-specific dorsal and ventral forebrain patterning was achieved using commercially available STEMdiff™ dorsal and ventral forebrain organoid differentiation kits (STEMCELL Technologies). Briefly, feeder-free iPSCs were dissociated into a single-cell suspension and seeded into AggreWell™800 plates to generate uniform cellular aggregates. Aggregation and early neural induction were carried out from day 0 to day 6 in forebrain organoid formation medium, with daily partial medium changes to support uniform aggregate formation. On day 6, aggregates were harvested, filtered to remove single cells, and transferred to ultra-low attachment 6-well plates for expansion. From day 6 to day 25, organoids were cultured in forebrain organoid expansion medium, with the medium fully changed every two days, to maintain controlled aggregate density and prevent premature fusion. Dorsal forebrain organoids were patterned without additional supplements, whereas ventral identity was induced by adding proprietary ventralizing components provided by the manufacturer. From day 25 to day 43, organoids were transitioned to forebrain organoid differentiation medium to promote neuronal differentiation and early circuit formation. Starting on day 43, organoids were maintained long-term in neural organoid maintenance medium with medium changes every 2–3 days (Xu et al. 2025; Vasan et al. 2025). Organoids were cultured for up to eight months *in vitro* and characterized at defined developmental stages to assess regional identity, cellular composition, cortical-like organization, and functional maturation.

**Figure 1.**
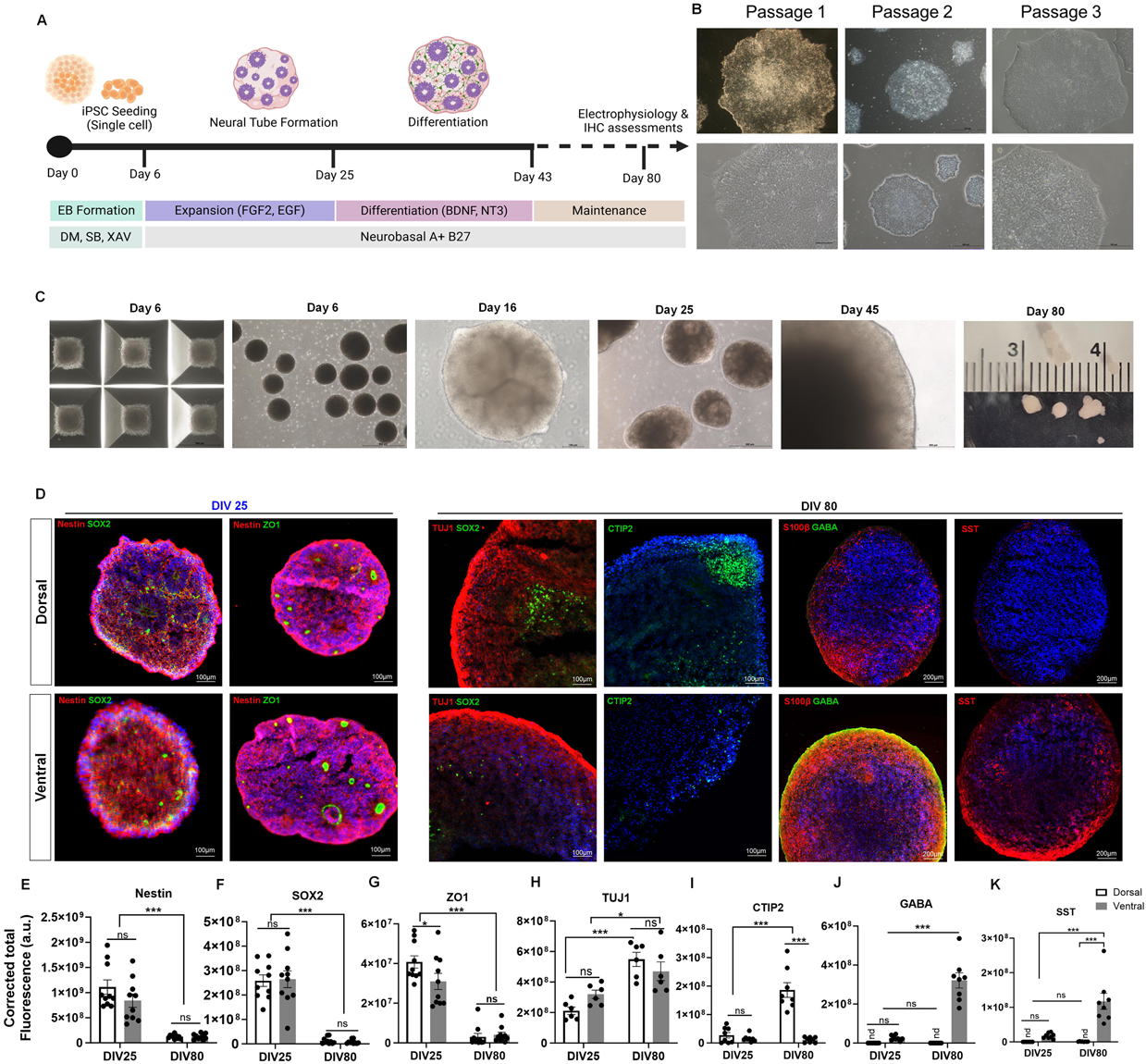
Generation and characterization of human forebrain organoids. **(A)** Schematic of the dual-SMAD inhibition protocol for the generation of dorsal and ventral human forebrain organoids (hFBOs). **(B)** Representative images of hiPSC colonies during expansion prior to differentiation. **(C)** Representative bright-field images illustrating hFBO development from embryoid body formation through progressive organoid growth and maturation The rightmost panel shows day 80 organoids alongside a ruler with centimeter markings for size reference. **(D)** Representative immunofluorescence images of dorsal and ventral hFBOs at DIV 25 and DIV 80 showing expression of developmental, progenitor, neuronal, and regional markers. **(E–K)** Quantification of fluorescence intensity for Nestin **(E)**, SOX2 **(F)**, ZO-1 **(G)**, TUJ1 **(H)**, CTIP2 **(I)**, GABA **(J)**, and SST **(K)** in dorsal and ventral hFBOs at DIV 25 and DIV 80. Data are presented as mean ± SEM (n = 10 hFBOs per group from 3 independent batches across 2 hiPSC lines.) and were analyzed by two-way ANOVA with DIV and regional identity as factors, followed by Tukey’s multiple-comparisons post hoc test. Fluorescence intensity is reported as corrected total cell fluorescence (CTCF; a.u.).

### Histological Preparation

On designated sampling days, organoids were randomly selected using a cut-end P1000 pipette tip and washed twice in phosphate-buffered saline (PBS). Organoids were then fixed in 4% paraformaldehyde (PFA) overnight at 4 °C. Following fixation, organoids were washed twice with PBS and cryoprotected by incubation in 30% sucrose (Acros Organics) at 4 °C until fully submerged. Organoids were subsequently embedded in optimal cutting temperature (OCT) compound (Tissue-Tek, Sakura), snap-frozen on dry ice, and stored at −80 °C or −20 °C until sectioning. Cryosections (20 µm thickness) were cut using a cryostat (Leica CM 1950), mounted onto Superfrost Plus slides (Fisher Scientific), and stored at −80 °C or −20 °C until further processing (McCrimmon et al. 2025).

### Immunofluorescence Staining

Cryosectioned slides were air-dried at room temperature for 15 minutes and washed three times in PBS (10 minutes per wash) to remove OCT and rehydrate the tissue. Heat-mediated antigen retrieval was performed by immersing slides in 1× Target Retrieval Solution (Dako) and steaming for 20 minutes. Slides were then washed with 0.05% PBST (phosphate-buffered saline containing Tween-20; Sigma) and permeabilized and blocked in 0.5% PBS–Triton X-100 (Sigma) supplemented with 5% donkey or goat serum (Dako) for 1 hour at room temperature. Primary antibodies (Supplementary Table 1) diluted in PBST containing 5% serum were applied overnight at room temperature. Following three washes in PBST (10 minutes per wash), slides were incubated with appropriate secondary antibodies (Supplementary Table 1), diluted in PBST with 5% serum, for 1 hour at room temperature. After three additional PBST washes, nuclei were counterstained with Hoechst 33342 (Sigma; 1:1,000 dilution) (Simorgh et al. 2023). Sections were mounted on glass slides using Vectashield mounting medium and imaged using a Zeiss Axioscope 5 microscope equipped with Stereo Investigator software (MBF Bioscience) and processed using ImageJ/Fiji software. Fluorescence intensity was quantified using identical imaging parameters across groups, with signals measured in standardized regions of interest and corrected for background fluorescence (data are presented as mean ± SEM; unpaired, two-tailed t-tests).

To assess apoptotic cell death in hFBOs, samples were processed for antigen retrieval and subjected to the Click-iT™ Plus Terminal deoxynucleotidyl transferase dUTP nick-end labeling (TUNEL) assay for in situ apoptosis detection (Thermo Fisher Scientific, C10617) according to the manufacturer’s instructions. Following completion of the TUNEL reaction, nuclei were counterstained with DAPI, and organoids were imaged by confocal microscopy. TUNEL-positive cells were identified by colocalization of nuclear and TUNEL fluorescence signals. For each developmental time point, 3–5 organoids per condition were analyzed.

### Electrophysiological assessments

Patch-clamp recordings were obtained in both current-clamp and voltage-clamp configurations at 80 and 150 DIV. For electrophysiological recordings, hFBOs were removed from culture medium and immediately embedded in 2% low–melting-point agarose (Sigma-Aldrich, A0576) prepared in PBS (in mM: 137 NaCl, 2.7 KCl, 10 Na₂PO₄, 1.8 KH₂PO₄; pH 7.4) (Simorgh et al. 2023). Dissolved agarose was maintained at 40 °C until use, and embedding was performed on ice to allow rapid gel solidification. Agarose-embedded organoids were sectioned using a Leica VT1200S vibratome in ice-cold, oxygenated (95% O₂ / 5% CO₂) sucrose-based artificial cerebrospinal fluid (aCSF) containing (in mM): 85 NaCl, 75 sucrose, 24 NaHCO₃, 25 glucose, 4 MgCl₂, 2.5 KCl, 1.25 NaH₂PO₄, and 0.5 CaCl₂. Slices were incubated at 32 °C for 20 minutes in an interface-holding chamber containing a 1:1 mixture of sucrose aCSF and recording aCSF composed of (in mM): 126 NaCl, 2.5 KCl, 2 CaCl₂, 2 MgCl₂, 1.25 NaH₂PO₄, 26 NaHCO₃, and 10 D-glucose. For voltage-clamp recordings of synaptic currents at 80 DIV, we used a modified aCSF containing an elevated KCl concentration (10 mM instead of 2.5 mM) to increase neuronal excitability and facilitate the detection of synaptic activity in neurons from hFBO at 80 DIV, which exhibited limited spontaneous activity under baseline conditions. All solutions were continuously bubbled with 95% O₂ / 5% CO₂ and maintained at pH 7.4. Slices were allowed to equilibrate for at least 1 hour at room temperature prior to recording (Afrasiabi et al. 2022; Subramanian et al. 2026; Gupta et al. 2012). Whole-cell recordings were obtained at 33–34 °C under infrared differential interference contrast (IR-DIC) optics using a Nikon Eclipse FN1 microscope equipped with a 40× water-immersion objective. Patch electrodes (3–6 MΩ) were fabricated using a dual-stage micropipette puller (Narishige PC-10). Signals were recorded using a MultiClamp 700B amplifier, digitized with a Digidata 1440A interface, and acquired using pClamp 10 software (Molecular Devices) at a sampling rate of 10 kHz. Spontaneous excitatory postsynaptic currents (sEPSCs) were recorded in voltage-clamp mode at a holding potential of −70 mV using a cesium-based internal solution containing (in mM): 140 cesium methanesulfonate, 5 NaCl, 10 HEPES, 0.2 EGTA, 2 MgATP, 0.2 Na₂GTP, and 5 QX-314 (pH 7.25; 270–285 mOsm) (Dovek et al. 2025). Current–voltage (I–V) relationships were generated using a step protocol ranging from −80 to +20 mV in 10 mV increments (200 ms duration). For current-clamp recordings, electrodes were filled with a potassium-based internal solution containing (in mM): 135 K-gluconate, 4 KCl, 10 HEPES, 4 MgATP, 0.3 Na₂GTP, and 10 phosphocreatine. Biocytin (0.2%) was included for post hoc morphological visualization of dendrites. Resting membrane potential (RMP) was measured immediately after break-in with zero holding current, and neurons were held at −70 mV during current-voltage (I/V) recordings. Following recordings, slices were fixed in 4% paraformaldehyde for subsequent histological analysis. Electrophysiological analyses, including intrinsic membrane properties and synaptic event quantification, were performed using EasyElectrophysiology software v2.6.3 (Easy Electrophysiology Ltd) (Dovek et al. 2025; Subramanian et al. 2026). Electrophysiological experiments were performed using dorsal and ventral hFBOs generated from two independent hiPSC lines across three independent differentiation batches.

### RNA Sequencing and Transcriptomic Analysis

For each sample, approximately 4–5 hFBOs were pooled at defined experimental time points. Total RNA was extracted using a column-based purification method according to the manufacturer’s instructions (PureLink™ RNA Mini Kit, Invitrogen) and on-column DNase treatment to remove genomic DNA contamination. RNA concentration and purity were assessed spectrophotometrically, and RNA integrity was evaluated using an RNA TapeStation (Agilent) prior to library preparation. All RNA samples had RNA integrity numbers (RIN) greater than 9, indicating high RNA quality. Sequencing libraries were prepared from poly(A)-enriched RNA using the NEBNext® Ultra™ II Directional RNA Library Prep Kit for Illumina and sequenced on an Illumina platform to generate paired-end reads. Raw sequencing reads were initially assessed for quality using FastQC (v.0.12.1) within Galaxy to evaluate per-base sequence quality, GC content, sequence duplication levels, and adapter contamination. Adapter sequences and low-quality bases were removed using Trimmomatic (v.0.39) with standard trimming parameters. Following trimming, read quality was reassessed using FastQC to confirm successful removal of adapters and low-quality sequences. High-quality reads were aligned to the human reference genome GRCh38/hg38 using HISAT2 (v.2.2.3), a splice-aware aligner optimized for transcriptomic data. Gene-level reading counts were generated from aligned BAM files using featureCounts (v.2.1.1) with the appropriate human gene annotation file (gencode v.47), and only uniquely mapped reads were retained for downstream analyses. Differential gene expression analysis was performed in R (v.4.5) using the DESeq2 (v1.50.2) package within Bioconductor 3.22. Raw gene count data were imported into DESeq2, normalized for sequencing depth, and analyzed using a negative binomial generalized linear model. DEGs were identified based on an adjusted p-value of below or equal to 0.05 and log_2_ fold changes <-1 and >1. Functional interpretation of differentially expressed genes was performed using Gene Ontology (GO) enrichment analysis with clusterProfiler (v4.16.4). Over-representation analysis was conducted using a hypergeometric test, and p-values were adjusted for multiple comparisons using the Benjamini–Hochberg method (Benjamini and Hochberg 1995). GO enrichment analyses were used to identify biological processes associated with neuronal identity, inflammatory signaling, and cellular stress responses, with enriched terms visualized using bar and dot plots (Among, Clusters, and Yu 2012; Liao, Smyth, and Shi 2014; Love, Huber, and Anders 2014; D. Kim et al. 2020; Bolger, Lohse, and Usadel 2014; Leggett et al. 2013). The Venn diagram was made with InteractiVenn 2.0 (Heberle et al. 2015). Underlying network interactions enriched within the DEGs were then queried at the protein level using STRING analysis (Szklarczyk et al. 2025) run against the homo sapiens NCBI taxonomy database (Id: 9606). Data were filtered for protein coding transcripts only, omitting antisense, long non-coding RNAs and microRNAs as well as divergent transcripts. Full networks (functional and physical interactions) were built using a high confidence (0.7) setting, an FDR of 5% and enriched clusters were then identified using the Leiden algorithm with a resolution parameter of 1. Edge identification considered evidence from textmining, experiments, databases, co-expression, neighborhood, gene fusion, co-occurrence and evidence transfer. The maximum number of interactors was set to query proteins only. Only clusters greater than 2 nodes or with statistical enrichment were considered for further analysis. A heatmap of the DEGs encoding the clustered proteins was made with pheatmap v.1.0.12 in R.

### Concussive Mechanical Injury Model in hFBOs

Mechanical injury to hFBOs was induced using the Cell Injury Controller II (CICII) (Fig 5A). Dorsal and ventral hFBOs were fully submerged in culture medium and positioned on an inflexible base within a multi-well plate for injury induction. The CICII regulates both the duration and magnitude of the applied pressure via a gas delivery system connected through a rubber tube to a circular well-adaptor plug that attaches to the top of the well to create an airtight chamber. A brief concussive pressure pulse (50 ms) was delivered, with the applied pressure continuously monitored using an in-line pressure transducer (Figure 5A) (Shaughness, Mike 2021). Injury severity was systematically varied by adjusting the peak pressure, with 1.5 psi defined as mild, 2.5 psi as moderate, and 3.5 psi as severe injury (Table 1). Following injury, organoids were returned to standard culture conditions and allowed to recover for various time points before histological and functional analyses.

**Table 1.**
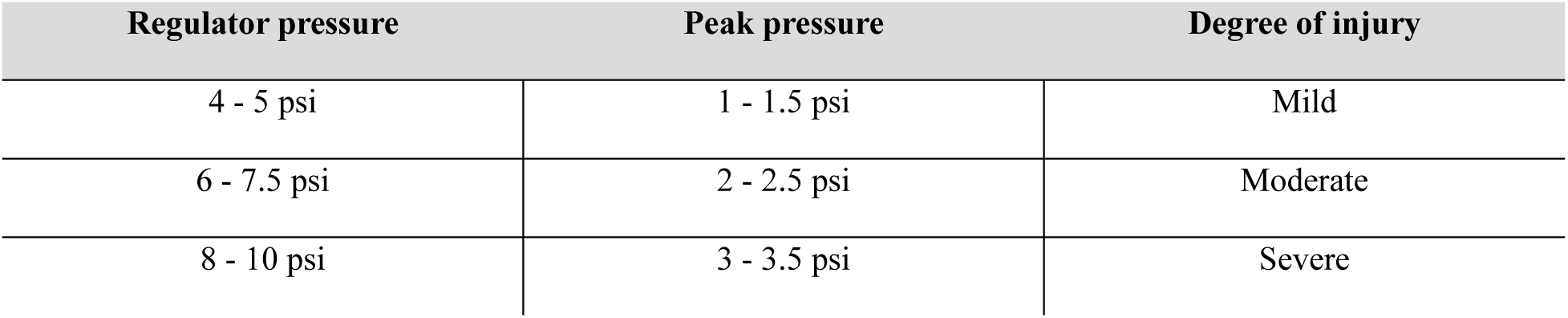
CICII injury severity.

### Calcium Imaging

Calcium dynamics in hFBOs were assessed at 150 DIV using the fluorescent calcium indicator Fluo-8 AM (Calcium Flux Assay Kit, Fluo-8 No-Wash, abcam). A stock solution was prepared by dissolving 50 µg of lyophilized Fluo-8 AM in 10 µL of sterile dimethyl sulfoxide (Sigma-Aldrich), yielding a final concentration of 5 mM. For staining, the stock solution was diluted to a working solution containing 5 µM Fluo-8 AM and 0.04% (v/v) Pluronic F-127 to enhance dye dispersion and cellular uptake. Before imaging, culture medium was aspirated from the wells containing hFBOs and replaced with the Fluo-8 AM working solution. Samples were incubated at 37 °C in a humidified 5% CO₂ incubator for 30 minutes to allow dye loading and intracellular de-esterification. Following incubation, the dye solution was replaced with Hanks’ Balanced Salt Solution (HBSS; Gibco) for imaging. Calcium imaging was performed using a Zeiss LSM880 inverted confocal microscope equipped with an FITC filter set (excitation: 490 nm; emission: 525 nm; cutoff: 515 nm). Time-lapse recordings were acquired to capture dynamic calcium transients over 120 frames at 1.26-second intervals, resulting in a total recording duration of approximately 150 seconds. Images were exported in CZI, TIFF, and MOV formats for downstream analysis and visualization. Post-acquisition analysis was performed using ImageJ. A whole-organoid ROI generated a global mean gray value (MGV) trace normalized to a pre-wave baseline. To capture wave-like propagation, three concentric rings (center, middle, rim) were defined; for each ROI, baseline fluorescence (F₀) was calculated from a quiescent pre-wave window, and signals were expressed as ΔF/F₀ using the equation ΔF/F₀ = (Fₜ − F₀) / F₀ (Okoro et al. 2026).

### Statistical analysis

All statistical analyses were performed using GraphPad Prism (version 10). Two healthy control human iPSC lines (SCTi003-A, female, and SCTi004-A, male) were used, with three independent organoid differentiation batches generated from each cell line. Multiple organoids were collected from each batch, and multiple sections or regions of interest (ROIs) were analyzed where applicable. Immunohistochemistry (IHC) images were analyzed by a researcher blinded to experimental conditions. Normality of data distributions was assessed using the D’Agostino– Pearson omnibus normality test, where sample size permitted. For IHC experiments, differences across developmental stages (25, 80, and 150 DIV) and regional identity (dorsal versus ventral hFBOs) were analyzed using two-way ANOVA, where appropriate. Similarly, intrinsic active and passive membrane properties were analyzed using two-way ANOVA. Spontaneous EPSCs were detected using the EasyElectrophysiology threshold-search algorithm, and all detected events were manually reviewed to exclude noise and artifacts that spuriously met detection thresholds. Synaptic activity was analyzed using cumulative probability distributions, with group differences assessed using the unpaired Kolmogorov–Smirnov test. Cumulative probability plots in Figure 3 were generated using an equal number of consecutive events from each cell. For calcium imaging experiments, comparisons among intact, mild, moderate, and severe injury groups were performed using one-way ANOVA followed by Tukey’s post hoc multiple-comparisons test. Where appropriate, additional post hoc tests were applied as specified in the corresponding figure legends. To account for the hierarchical structure of the data, measurements obtained from multiple sections or ROIs within the same organoid were treated as technical replicates rather than independent biological samples. Independent organoids and differentiation batches were identified separately, and the experimental unit used for each statistical analysis is specified in the corresponding figure legend. The number of biological replicates, technical replicates, and independent differentiation batches is reported for each experiment. Images or recordings that did not meet predefined quality-control criteria were excluded from analysis, where applicable. Statistical significance was set at p < 0.05. Data are presented as mean ± standard error of the mean (SEM), unless otherwise indicated. The exact statistical tests, sample sizes, and definitions of biological replicates are provided in the corresponding figure legends.

## Results

### Regional and Developmental Specification of Human Forebrain Organoids

TBI disrupts adult neurogenesis and alters excitatory-inhibitory circuit balance, contributing to long-term cognitive dysfunction and enhanced network excitability (Ngwenya and Danzer 2019; Neuberger, Swietek, et al. 2017). While rodent models have provided important insights, species-specific differences in human cortical development and interneuron maturation limit translational relevance. To address this gap, we sought to establish regionally-specified hFBO models that recapitulate dorsal (Glutamatergic) and ventral (GABAergic) developmental programs (Birey et al. 2017), enabling investigation of injury-induced alterations in neurogenesis and network function. We generated hFBOs using the dual SMAD-inhibition protocol adapted from Sloan et al. (Fig. 1A) (Sloan, Andersen, Pașca, et al. 2018). Human induced pluripotent stem cells (hiPSCs, female line, StemCell Technologies) were cultured under feeder-free conditions and passaged three times prior to differentiation. At the third passage, colonies exhibited compact, round morphology typical of healthy pluripotent stem cells, with clear colony borders and homogenous cells within that exhibited a high nuclear-to-cytoplasmic ratio (Fig. 1B). By day 6, aggregation of hiPSCs in low-adhesion multiwell plates resulted in relatively uniform embryoid bodies (EBs) that were transferred to ultra-low-attachment wells to facilitate neural induction (Fig. 1C). By day 16, the forming organoids displayed distinct neuroepithelial rosettes, indicating establishment of early radial organization (Finger et al. 2023). The organoids continued to expand in size, reaching approximately 2 mm in diameter by day 80, suggesting sustained growth and tissue maturation (Fig. 1C). Immunostaining confirmed that cellular identity progressed in a developmentally consistent manner across three independent batches of hFBOs, consistent with previously established forebrain organoid protocols (Sloan, Andersen, Pașca, et al. 2018). Dorsal and ventral regional identities were detected in all three batches. By 25 DIV, organoids consistently retained Nestin-expressing neural progenitor populations and organized radial architecture (Fig. 1D, left panel). At this stage, hFBOs recapitulated early neurodevelopmental features, as evidenced by strong expression of neural stem cell (NSC) and progenitor markers and the presence of organized neuroepithelial rosette-like structures. Immunohistological analysis of Nestin and SOX2, markers of radial glial neural progenitor cells, revealed multiple ventricular zone (VZ)–like architectures within both dorsal and ventral hFBOs. These cellular structures also expressed the tight junction marker ZO1, indicating that the hFBOs acquired epithelial identity (Fig. 1D). Quantitative analysis of immunostaining further confirmed high expression levels of these markers at this developmental stage, with no difference in Nestin and SOX2 expression between dorsal and ventral hFBOs (Fig. 1D, E–F). Interestingly, ZO1 expression was lower in ventral than in dorsal hFBO (Fig 1D, G). Immunostaining at 25 DIV revealed early but limited expression of CTIP2 and SST in hFBOs. However, CTIP2 and SST expression did not differ between dorsal and ventral hFBOs at this early developmental stage (Fig. 1I, K and supplementary 1B), suggesting that cell types were not specified at this point.

To determine the persistence of neurogenesis and maintenance of progenitor populations, we examined the expression of markers of neural progenitors at 80 DIV. Nestin-, SOX2-, and ZO1-positive cells persisted in both dorsal and ventral hFBOs at 80 DIV, with no significant differences between the two regions. However, the proportion of Nestin-, SOX2-, and ZO1-positive cells was significantly reduced compared with the earlier time point at 25 DIV, indicating a decline in progenitor populations as the organoids matured and differentiated (Fig. 1E-G, and Supplementary 1A). In contrast, at 80 DIV, neuronal differentiation was evident (Fig. 1D, panel right), as demonstrated by strong expression of the neuronal marker βIII-tubulin (TUJ1) in both dorsal and ventral hFBOs (Fig 1 D, H, Supplemental Fig 1A). Clear subtype specifications also emerged at this stage. CTIP2-expressing glutamatergic cortical neurons were preferentially enriched in dorsal hFBOs, whereas GABAergic interneurons, including somatostatin (SST)-positive cells, were enriched in ventral hFBOs, demonstrating region-specific neuronal identity consistent with forebrain patterning (Fig. 1D). In addition, expression of the glial marker S100β was observed in both dorsal and ventral hFBOs (Fig. 1D). Quantitative analysis revealed differential enrichment of neuronal subtype markers for gluatamatergic and GABAergic in their respective dorsal and ventral regions (Fig. 1I-K).

We further extended the developmental profile and examined hFBOs at 150 DIV, when parvalbumin (PV)-expressing interneurons, a major inhibitory cell type, have been shown to develop (Varela et al. 2025). At 150 DIV, ventral (but not dorsal) hFBOs showed neuronal expression of PV, consistent with progressive maturation and region-specific cellular patterning (Walsh et al. 2024) (Supplementary Fig. 1C). Together, these findings establish a developmental framework for cell type specification in dorsal and ventral hFBO in our hands. At 25 DIV, organoids are enriched in neural progenitor populations and exhibit ventricular zone-like architecture, reflecting early neurodevelopmental stages. By contrast, at 80 DIV, hFBOs display robust neuronal differentiation and the emergence of region-specific neuronal subtypes. This staged progression highlights the transition from progenitor-rich structures to more mature neuronal populations, providing a framework to study early developmental processes and later neuronal specialization in a hFBO model.

### Early Dorso-Ventral Forebrain Patterning Drives Distinct Neurogenic and Synaptic Transcriptional Programs in hFBOs

Because immunohistochemical analyses of hFBOs at 25 DIV revealed no differences in cellular composition, except a small difference in ZO1 expression between dorsal and ventral organoids, we examined whether transcriptional profiling could uncover region-specific molecular signatures at this early stage. Accordingly, bulk RNA sequencing was performed on hFBOs at 25 DIV. To assess regional transcriptional identity, we directly compare dorsal and ventral organoids. PCA plots showed that dorsal and ventral organoids segregated into two distinct clusters (Fig. 2A). Based on a p.adjust cut-off of 0.05 and a log_2_ threshold of 1 and -1, respectively, 197 transcripts were identified as specific to ventral hFBOs and 350 as specific to dorsal identity (Fig. 2B). Further graphing of the differentially expressed genes in the form of volcano plots revealed robust differential expression of genes associated with excitatory and inhibitory neuronal lineages in either DEG set, consistent with expected dorsal-ventral patterning (Fig. 2C). Gene Ontology (GO) enrichment analysis of all differentially expressed genes revealed significant overrepresentation of biological processes associated with forebrain development, pattern specification, axonogenesis, regulation of neural precursor cell proliferation, and neuron migration. Additional enrichment for synapse assembly and regulation of glutamatergic synaptic transmission indicated that early circuit-related programs are already being established at 25 DIV (Fig. 2D). Molecular function categories further supported active developmental signaling, including receptor ligand activity, growth factor activity, G protein–coupled receptor activity, glutamate receptor activity, and DNA-binding transcription factor activity. Consistent with these findings, cellular component analysis demonstrated enrichment of genes associated with synaptic membranes, receptor complexes, axon termini, and both glutamatergic and GABAergic synaptic structures (Fig. 2D), validating the identity of cells within organoids as forebrain specific. To characterize regional specification in the organoids, a protein-protein-interaction (PPI) analysis was next carried out using STRING software (Szklarczyk et al. 2025) that assumed the translation of identified DEGs into functional proteins and considered their likelihood for potential interaction. The ventral PPI network encompassed 179 nodes and 42 interactions (edges), with an average local clustering coefficient of 0.0355 and enrichment of p < 2.26e-13 (average node degree 0.469). The dorsal PPI network encompassed 307 nodes and 139 interactions (edges), with an average local clustering coefficient of 0.08 and enrichment of p<9.69e-51 (average node degree 0.906). Based on their centrality and interactions (≥ 3 node degrees), the proteins highlighted as important nodes are shown in color in Fig. 2E and Fig. 2F, respectively. Consistent with ventral identity, the ventral hFBO network analysis identified clusters enriched in functions associated with GABAergic neuron differentiation and ventral midline development, highlighting the importance of the nodes SHH and NKX2.1. These were not only identified as network hubs but also represent the 11^th^ and 19^th^ highest regulated transcripts in the ventral hFBO organoids (log_2_FC of -3.6 and -3.09, respectively) (see also Fig. 2C). Overall, of the 30 topmost regulated transcripts in the ventral hFBOs, 10 were represented in the network clusters with log_2_FC values ranging from -3.78 to -2.86 (*SERPINF2, TFF3, SHH, CER1, FOXA1, NKX6.1, NKX2.1, GATA3, PITX2 and NKX6.2*) (Fig. 2G) with most of these having been reported to be expressed in ventral forebrain (Zhao et al., 2014; Craven et al., 2004; Nat et al., 2012).

**Figure 2.**
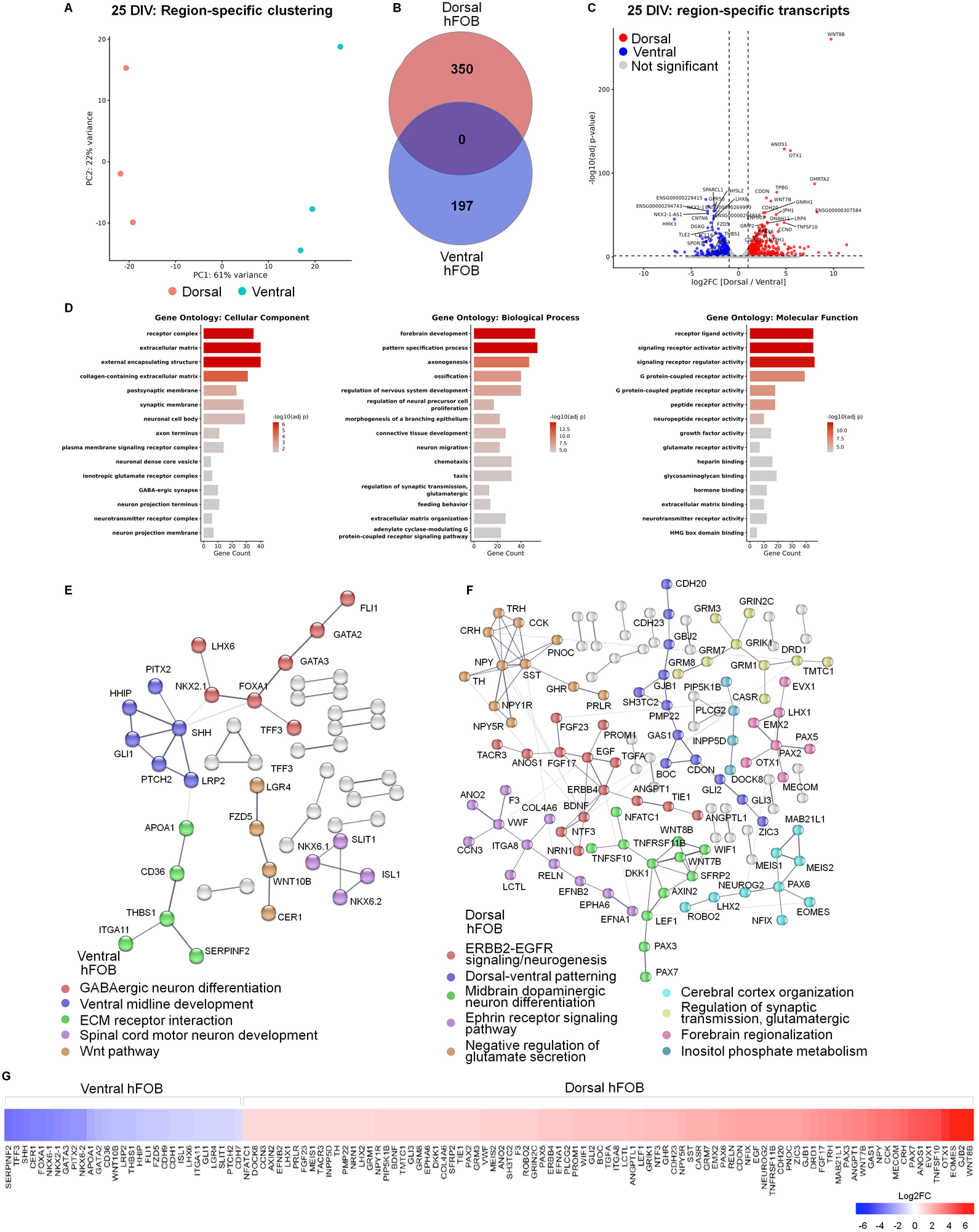
Establishment of dorsoventral transcriptional identity in early hFBOs. **(A)** Principal component and unsupervised clustering analyses revealed clear segregation of dorsal and ventral organoids, indicating robust and reproducible regional transcriptional patterning. **(B)** Venn diagram separating the down-regulated DEGs representing the ventral hFBO-specific transcripts from the up-regulated DEGs that correspond to the dorsal-specific transcripts. **(C)** Differential expression results were visualized using a volcano plot displaying the log₂ fold change versus the negative log10 of the adjusted p-value, with grey dots representing the non-significantly regulated transcripts (p.adjust >0.05, log_2_FC -1-1). **(D)** Gene Ontology enrichment analysis across all DEGs highlights distinct functional programs, with dorsal organoids enriched for neurodevelopmental, axonogenesis, and glutamatergic synaptic pathways, and ventral organoids enriched for interneuron specification, migration, and GABAergic synaptic processes. (D, E) STRING protein-protein interaction analysis using ventral (D) and dorsal (E)-specific DEGs to chart region-specifically enriched network clusters at the protein level. Nodes correspond to differentially abundant proteins represented by their gene symbol, while lines between nodes refer to edges that represent interaction confidence. Clusters <3 nodes or with statistically insignificant enrichment were excluded from further analysis and are shown in light gray. All disconnected nodes are hidden. **(G)** Heatmap visualizing transcripts encoding proteins found in the cluster enrichment based on log_2_FC.

Similarly, PPI analysis on dorsal hFBOs confirmed their dorsal regionalization due to enrichment in glutamatergic synaptic transmission, cerebral cortex organization and dorsal-ventral patterning association (Fig. 2F), with interconnectivity between the glutamate secretion and glutamatergic transmission clusters, as well as ERBB and ephrin signaling. As such, ERBB4 is preferentially found in interneurons as they migrate to the dorsal telencephalon and those restricted to the frontal dorsal cortex (Yau et al. 2003; Neddens et al. 2011) suggesting involvement in regional brain development and connectivity. 15 of the 35 topmost regulated dorsal hFBO transcripts were included in this PPI enrichment (log2FC 3.25-8.85), indicating a strong correlation between the two analyses. While not included in the cluster interaction enrichment, the dorsal marker *DMRTA2* was the third highest regulated DEG in the dorsal hFBOs (log_2_FC=7.9) (Fig. 2B) again in agreement with established dorso–ventral transcriptional programs (Anirudhan et al. 2025; Shen et al. 2025) (Fig. 2G). Together, these results demonstrate that by 25 DIV, hFBO robust and regionally appropriate transcriptional programs supporting early neurogenesis, neuronal migration, and lineage specification. This stage therefore provides a biologically relevant window for investigating mechanisms governing excitatory and inhibitory neuronal development. The distinction between dorsal and ventral organoids is particularly important, as these regions give rise to excitatory projection of neurons and inhibitory interneurons, respectively, enabling region-specific analyses of neurogenic trajectories.

### Temporal Maturation of Functional Excitatory Networks in Human Forebrain Organoids

Given the emergence of neuronal markers at 80 DIV (TUJ1, CTIP2, GABA, SST), we next asked whether these molecular signatures were accompanied by functional maturation. Whole-cell patch-clamp recordings were obtained from neurons in dorsal and ventral hFBOs at 80 and 150 DIV (Fig. 3). During whole-cell recordings, neurons were filled with biocytin for post hoc visualization of the neurites (Fig. 3A, B). Neurons at 80 DIV displayed shorter dendritic processes and reduced arbor complexity compared with neurons at 150 DIV (Fig. 3A, B). At 80 DIV, neurons in both dorsal and ventral hFBOs generated rudimentary wide action potentials in response to depolarizing current injections (Fig. 3C), indicative of emerging but still immature cellular excitability. By contrast, neurons recorded at 150 DIV exhibited large-amplitude repetitive action potentials with short half-width (Fig. 3D), reflecting greater physiological maturation. We analyzed passive membrane properties including resting membrane potential (RMP), input resistance (R_in_), and active firing properties such as action potential threshold, amplitude, and half-width in both dorsal and ventral hFBOs at 80 and 150 DIV (Fig. 3E-I).

**Figure 3.**
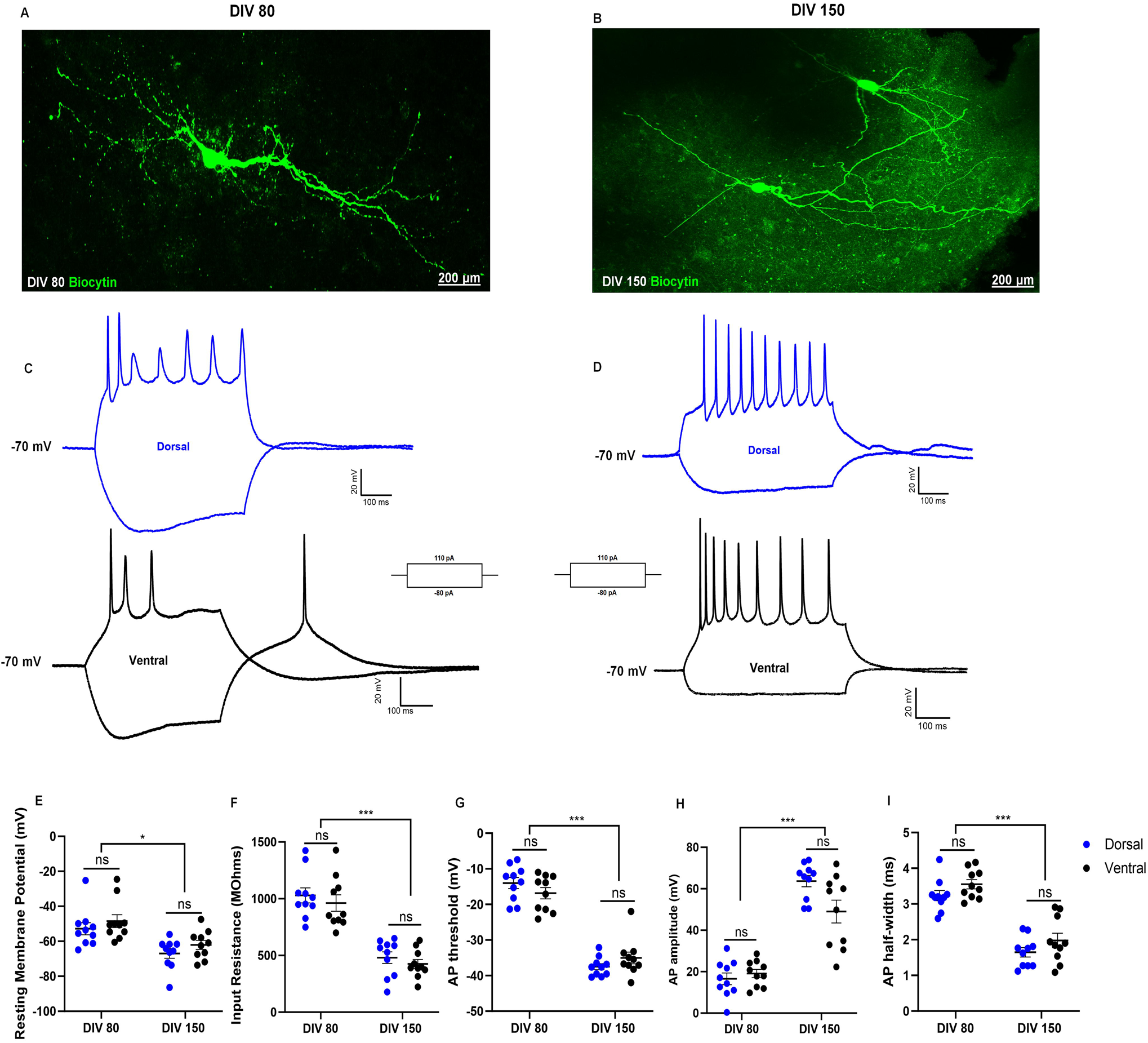
Functional maturation of human forebrain organoids. **(A, B)** Representative images of biocytin-filled neurons recorded from human forebrain organoids (hFBOs) at 80 DIV **(A)** and 150 DIV **(B). (C, D)** Representative current-clamp recordings from dorsal (above) and ventral (below) hFBO neurons at 80 DIV **(C)** and 150 DIV **(D)** in response to hyperpolarizing (−80 pA) and depolarizing (+110 pA) current injections from a holding potential of −70 mV.**(E-I)** Summary plots of resting membrane potential (E), input resistance (F), action potential threshold (G), action potential amplitude (H) and action potential half-width (I) in dorsal and ventral hFBOs at 80 and 150 DIV. Data are presented as mean ± SEM (n = 10 cells per group from 3 independent batches derived from 2 hiPSC lines) and were analyzed by two-way ANOVA with DIV and regional identity as factors, followed by Tukey’s multiple-comparisons post hoc test.

Analysis of the RMP identified a significant effect of hFBO DIV (F (1,36) = 19.51, p < 0.0001), with no effect of regional specification (F (1,36) = 2.199, p = 0.1468) (Fig. 3F). In dorsal hFBOs, RMP became more hyperpolarized from −52.82 ± 3.48 mV at 80 DIV to −66.96 ± 2.72 mV at 150 DIV. Similarly, in ventral hFBOs, RMP shifted from −48.43 ± 3.67 mV at 80 DIV to −62.03 ± 2.55 mV at 150 DIV. No significant differences were observed between dorsal and ventral hFBOs at either 80 DIV (p = 0.7568) or 150 DIV (p = 0.6865; two-way ANOVA followed by Tukey’s post hoc test, n = 10 cells/group from 3 hFBOs, derived from 2 independent hiPSC lines) (Fig. 3F). Consistent with this developmental shift in RMP, membrane input resistance (R_in_) also changed with hFBO maturation. R_in_ showed a significant main effect of DIV (F (1,36) = 84.96, p < 0.0001), with no significant effect of regional specification (F (1,36) = 1.094, p = 0.3026) (Fig. 3F). In dorsal hFBOs, R_in_ significantly decreased from 1030 ± 66.47 MΩ at 80 DIV to 481.0 ± 51.42 MΩ at 150 DIV (p < 0.0001). Similarly, in ventral hFBOs, R_in_ decreased from 961.8 ± 72.39 MΩ at 80 DIV to 425.7 ± 39.28 MΩ at 150 DIV (p < 0.0001). No significant differences were observed between dorsal and ventral hFBOs at either 80 DIV (p = 0.8475) or 150 DIV (p = 0.9095; two-way ANOVA followed by Tukey’s post hoc test, n = 10 cells/group from 3 hFBOs/group, derived from 2 independent hiPSC lines) (Fig. 3F). Alongside these changes in passive membrane properties, active membrane properties also showed pronounced developmental changes. The action potential threshold showed a significant main effect of DIV (F (1,36) = 207.8, p < 0.0001), with no significant effect of regional specification (F (1,36) = 0.01827, p = 0.8932) (Fig. 3G). Neurons at 80 DIV exhibited a more depolarized action potential threshold compared to neurons at 150 DIV in both dorsal (threshold in mV, 80 DIV: −14.01 ± 1.55; dorsal: −37.49 ± 0.85, p<0.05 two-way ANOVA followed by Tukey’s post hoc test, n = 10 cells/group from 3 hFBOs/group, derived from 2 independent hiPSC lines) ventral hFBOs (threshold in mV, 80 DIV: −16.86 ± 1.58; 150 DIV: −35.03 ± 1.65 mV p < 0.05 by two-way ANOVA followed by Tukey’s post hoc test, n = 10 cells/group from 3 hFBOs/group, derived from 2 independent hiPSC lines), likely reflecting developmental changes in sodium channel density (Fig. 3G). This developmental maturation was further reflected in action potential amplitude. Action potential amplitude showed a significant main effect of DIV (F (1,36) = 157.8, p < 0.0001), with no significant effect of regional specification (F (1,36) = 1.747, p = 0.1946) (Fig. 3H). In dorsal hFBOs, action potential amplitude significantly increased from 16.53 ± 2.80 mV at 80 DIV to 63.70 ± 2.75 mV at 150 DIV (p < 0.0001). Similarly, in ventral hFBOs, amplitude increased from 19.10 ± 1.97 mV at 80 DIV to 52.64 ± 4.69 mV at 150 DIV (p < 0.0001). No significant differences were observed between dorsal and ventral hFBOs at either 80 DIV (p = 0.9418) or 150 DIV (p = 0.0886; two-way ANOVA followed by Tukey’s post hoc test, n = 10 cells/group from 3 hFBOs/group, derived from 2 independent hiPSC lines) (Fig. 3H). Consistent with these changes in action potential threshold and amplitude, action potential kinetics also differed with maturation. Similarly, action potential half-width showed a significant main effect of DIV (F (1,36) = 104.7, p < 0.0001) and a significant main effect of regional specification (F (1,36) = 4.419, p = 0.0426) (Fig. 3I). In dorsal hFBOs, action potential half-width decreased from 3.24 ± 0.14 ms at 80 DIV to 1.65 ± 0.13 ms at 150 DIV. Similarly, in ventral hFBOs, half-width decreased from 3.55 ± 0.13 ms at 80 DIV to 1.98 ± 0.20 ms at 150 DIV. Although a significant main effect of regional specification was observed, post hoc comparisons did not identify significant differences between dorsal and ventral hFBOs at either 80 DIV (p = 0.4752) or 150 DIV (p = 0.4371; two-way ANOVA followed by Tukey’s post hoc test, n = 10 cells/group from 3 hFBOs/group, derived from 2 independent hiPSC lines) (Fig. 3I). Half-width was significantly reduced from 80 to 150 DIV in both dorsal and ventral hFBOs (Fig. 3I). Together, these findings demonstrate temporal maturation of passive and active properties of neurons in hFBO. While neurons at 80 DIV exhibited immature electrophysiological characteristics, neurons at 150 DIV displayed more mature intrinsic membrane properties and robust firing activity. Notably, these properties were comparable between dorsal and ventral hFBOs, indicating that maturation of cell intrinsic physiology progresses similarly across both regional identities.

Having established that neurons appear functionally mature at 150 DIV, we next examined the development of synaptic function in hFBOs by recording spontaneous excitatory postsynaptic currents (sEPSCs) at a holding potential of −70 mV (Fig. 4). sEPSCs were recorded over 5–10 min from neurons in dorsal and ventral hFBOs. At 80 DIV, no spontaneous EPSCs were detected in either dorsal or ventral hFBOs in recordings from five cells obtained from three independent organoids per region (not shown). To evaluate synaptic competence at this stage, we increased the extracellular potassium to 10 mM (Hi-K) to enhance network activity. In the presence of Hi-K, neurons from hFBOs at 80 DIV exhibited EPSCs, confirming the presence of functional synapses and activity-dependent neurotransmission (Fig. 4A). At DIV 80, the EPSCs inter-event interval (IEI) in hi-K was shorter in dorsal compared to ventral hFBOs (IEI in seconds; Dorsal-Median: 0.167, IQR: 0.23; n= 4 cells from 3 hFBOs, Ventral-Median: 0.5, IQR: 1.4; n = 3 cells from 3 hFBOs across 3 independent batches derived from 2 hiPSC lines, *P*<0.0001 by K-S test, Cohen’s d = 0.67), indicating a higher frequency of glutamatergic synaptic events in dorsal hFBOs (Fig. 4B). EPSC amplitude in neurons from dorsal hFBOs was smaller than in ventral hFBOs (Amplitude in pA; Dorsal: Median: 15.98, IQR: 10.47; n= 4 cells from 3 hFBOs derived from 3 independent batches across 2 hiPSCs, Ventral: Median: 23.33, IQR: 33.01; n = 3 cells from 3 hFBOs derived from 3 independent batch across 2 hiPSCs, P<0.0001 by K-S test, Cohen’s d = 0.81) (Fig 4C). In contrast to the need for elevated potassium to evoke synaptic events in hFBOs at 80 DIV, spontaneous EPSCs were readily observed in neurons from both dorsal and ventral hFBOs at 150 DIV (Fig. 4D), indicating the emergence of spontaneous neuronal activity. At 150 DIV, sEPSCs in neurons from dorsal hFBOs exhibited shorter IEI than in ventral hFBOs (IEI in s; Dorsal: Median: 0.15, IQR:0.53; n = 6 cells from 3 hFBOs derived from 3 independent batches across 2 hiPSCs, Ventral: Median: 0.27, IQR: 0. 73, n = 4 cells from 3 hFBOs derived from 3 independent batches across 2 hiPSCs. *P*<0.0001 by K-S test, Cohen’s d = 0.16), indicating higher sEPSC frequency (Fig. 4E). Additionally, sEPSC amplitude was higher in dorsal relative to ventral hFBOs at 150 DIV (Amplitude in pA: Dorsal: Median: 17.7, IQR: 9.9, n = 6 cells from 3 hFBOs derived from 3 independent batches across 2 hiPSCs, Ventral: Median: 10.03, IQR: 8.94, n = 4 cells from 3 hFBOs derived from 3 independent batch across 2 hiPSCs, *P*<0.0001 by K-S test, Cohen’s d = 0.63) (Fig. 4F). Collectively, these findings demonstrate that by 150 DIV, hFBOs exhibit spontaneous activity and regionally distinct synaptic function, supporting this later stage as optimal for functional analyses.

**Figure 4.**
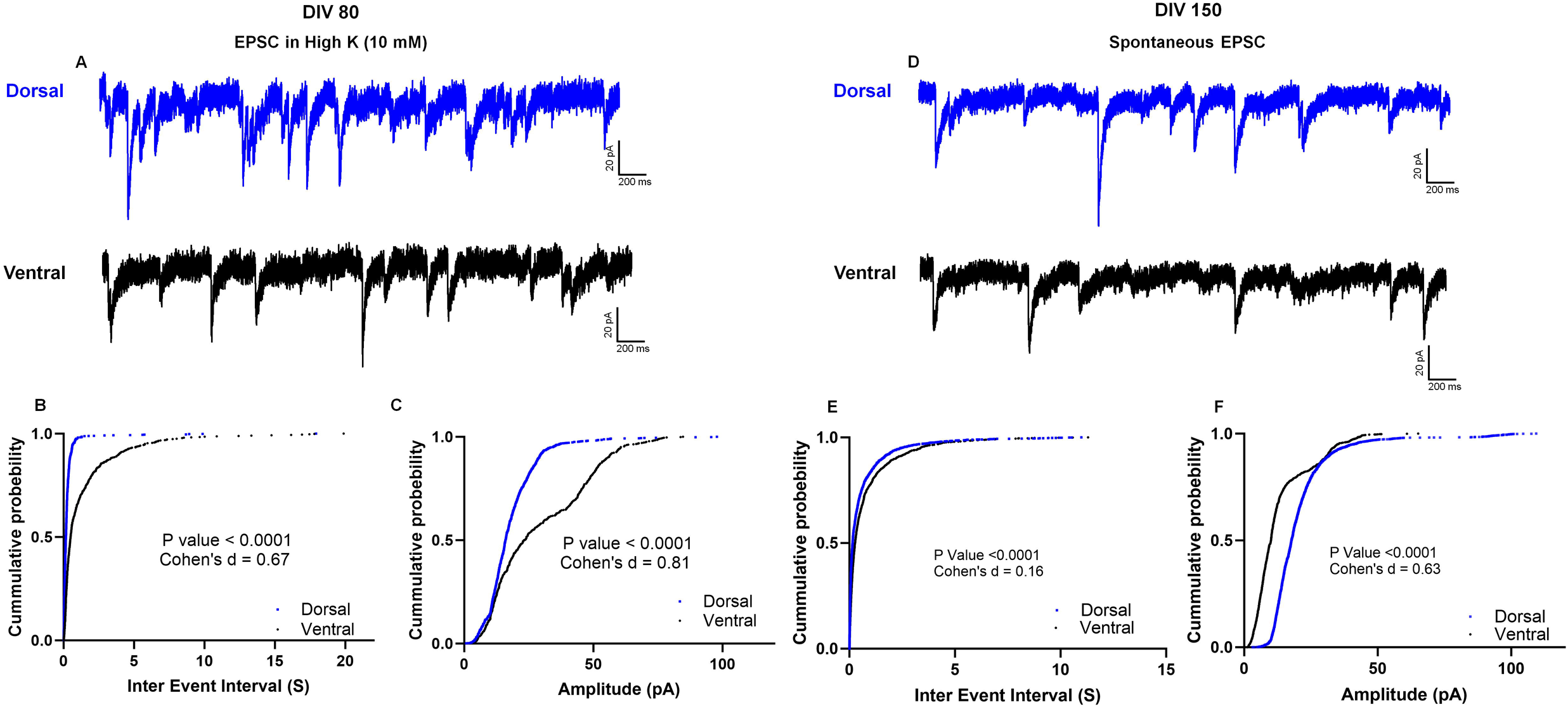
Maturation of excitatory synaptic activity in human forebrain organoids. **(A)** Representative voltage-clamp recordings illustrate EPSCs in neurons from dorsal and ventral hFBOs at 80 DIV in the presence of 10 mM extracellular K⁺. **(B)** Cumulative probability distribution of EPSC inter-event intervals (IEIs) at 80 DIV, showing shorter IEIs in dorsal compared with ventral hFBOs (n = 3 cells from 3 hFBOs across 3 independent batches derived from 2 hiPSC lines). **(C)** Cumulative probability distribution of EPSC amplitudes at 80 DIV, showing smaller amplitudes in dorsal compared with ventral hFBOs. Recordings were obtained from Dorsal: n = 4 cells from 3 hFBOs; ventral: n = 3 cells from 3 hFBOs, derived from 3 independent batches across 2 hiPSC lines. **(D)** Representative voltage-clamp recordings from dorsal and ventral hFBO neurons at 150 DIV showing spontaneous EPSCs (sEPSCs). **(E)** Cumulative probability distribution of sEPSC IEIs at 150 DIV, showing shorter IEIs in dorsal compared with ventral hFBOs. **(F)** Cumulative probability distribution of sEPSC amplitudes at 150 DIV, showing larger amplitudes in dorsal compared with ventral hFBOs. At 150 DIV, recordings were obtained from Dorsal: n = 6 cells from 3 hFBOs; ventral: n = 4 cells from 3 hFBOs, derived from 3 independent batches across 2 hiPSC lines. Cumulative probability distributions were compared using the unpaired Kolmogorov–Smirnov (K–S) test, with p values and effect size indicated in the figure.

### Injury Severity not Regional Specification influences Cell Death in a Novel Model of Concussive Trauma in Human Forebrain Organoids

To model concussive injury in a human-relevant 3D system, we adapted the CICII, originally designed for two-dimensional cultures, for use in hFBOs. Organoids were positioned in low-adhesion 6-well plates with a non-deformable base to prevent displacement during impact. To ensure an airtight seal and prevent gas leakage during pressure delivery, we engineered a custom adaptor with an O-ring gasket that securely connected the device nozzle to the well. Pressure parameters were calibrated to allow uniform mechanical force transmission. The system was optimized by calibrating pressure intensity, chamber volume, and organoid positioning to ensure uniform exposure to pressure while maintaining structural integrity (Fig. 5A; Table 1). This system enabled reproducible delivery of graded concussive injury in a 3D system. To determine the effect of injury severity on cell death, dorsal hFBOs at 25 DIV were subjected to graded pressure intensities corresponding to mild, moderate, and severe injury (Table 1) and examined at various time points following injury (Fig. 5B). Cell death was quantified at 1 and 3 days post-injury (dpi) by TUNEL staining as the percentage of TUNEL-positive cells relative to total DAPI-positive nuclei. The proportion of TUNEL-positive cells increased progressively with injury intensity, indicating greater cell death at higher injury pressures compared with intact controls (Fig. 5C–E). At 1 dpi, TUNEL-positive cells increased with injury intensity (Intact: 11.82 ± 1.31%; Mild PSI: 29.18 ± 2.65%; Moderate: 30.55 ± 2.57%; Severe: 46.59 ± 3.29%; mean ± SEM; n = 10 bins from 5 hFBOs per group, derived from 3 independent batches from one hiPSC line; One-way ANOVA with Tukey’s multiple-comparisons post hoc test) (Fig. 5C, D). Similarly, organoids examined at 3 dpi showed an increase in TUNEL-positive cells with injury intensity (Intact: 11.23 ± 1.38%; Mild: 16.76 ± 2.73%; Moderate: 31.94 ± 2.66%; Severe: 50.36 ± 3.33%; mean ± SEM; n = 10 bins from 5 hFBOs per group, derived from 3 independent batches from one hiPSC line; One-way ANOVA with Tukey’s multiple-comparisons post hoc test), further demonstrating an injury severity-dependent increase in cell death (Fig. 5E). To determine whether injury responses differed between regional identities, we subjected ventral hFBOs to increasing injury intensities (mild, moderate, and severe) and assessed cell death at 1 and 3 dpi (Supplementary Fig. 2A). TUNEL staining increased with injury intensity, indicating greater cell death following higher injury pressures compared with intact controls. At 1 dpi, TUNEL-positive cells progressively increased with injury intensity (Intact: 12.56 ± 1.08%; Mild: 27.98 ± 1.63%; Moderate: 37.66 ± 1.12%; Severe: 49.12 ± 2.45%; mean ± SEM; n = 8 bins from 3 hFBOs/group; One-way ANOVA with Tukey’s multiple-comparisons post hoc test) (Supplementary Fig. 2B, and C). At 3 dpi, TUNEL-positive cells increased with injury intensity (Intact: 13.82 ± 0.85%; Mild: 15.43 ± 0.92%; Moderate: 39.86 ± 1.13%; Severe: 54.31 ± 2.16%; mean ± SEM; n = 8 bins from 5 hFBOs per group, derived from 3 independent batches from one hiPSC line; One-way ANOVA with Tukey’s multiple-comparisons post hoc test) (Supplementary Fig. 2B, D). Notably, the Mild group did not differ significantly from intact controls, consistent with the response observed in dorsal hFBOs. Together, these findings demonstrate an injury severity-dependent increase in cell death, with comparable response patterns between ventral and dorsal hFBOs.

**Figure 5.**
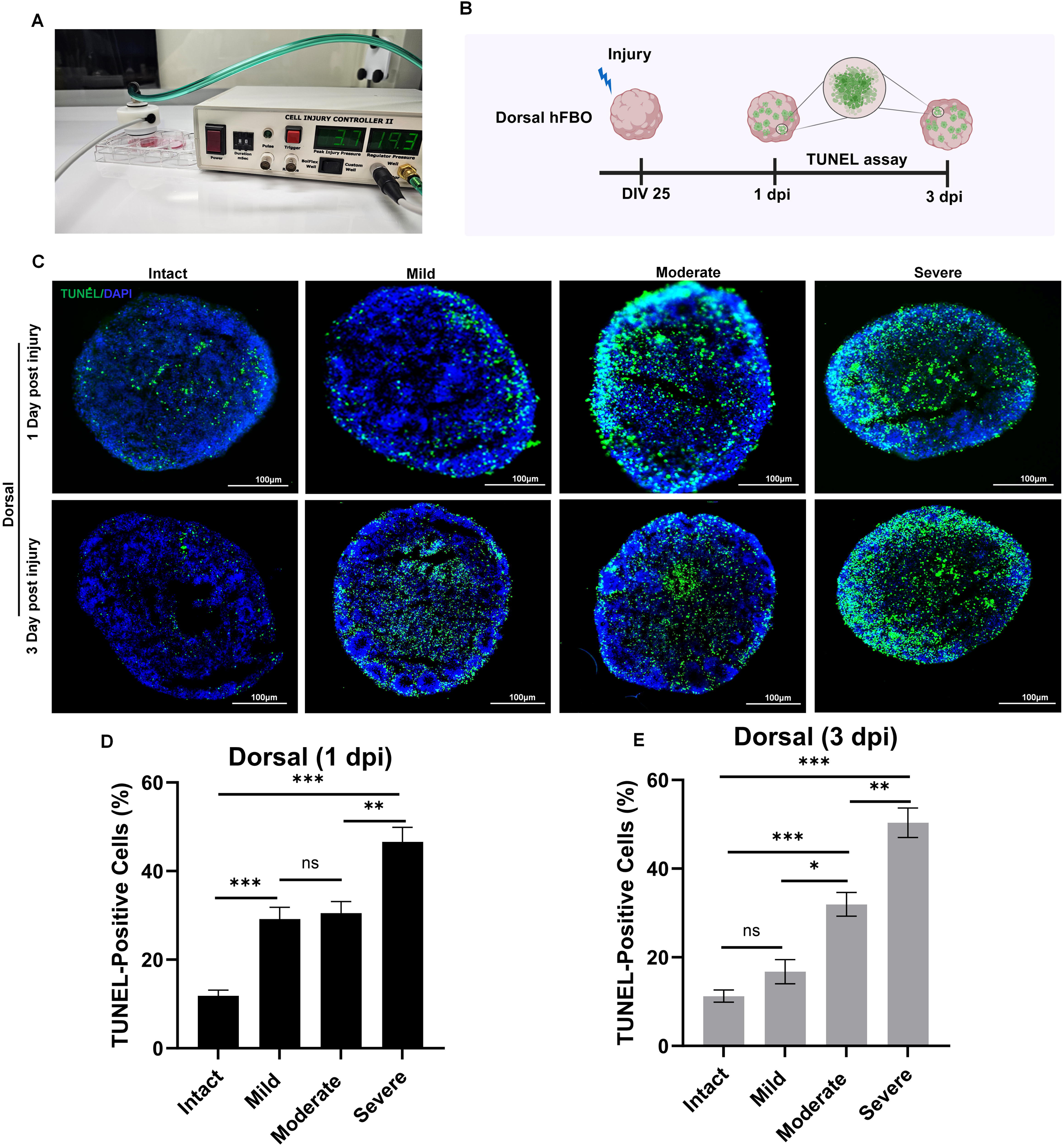
Injury severity-dependent cell death in dorsal human forebrain organoids following concussive injury. **(A)** Experimental setup used to deliver controlled mechanical impact to hFBOs. **(B)** Schematic of the experimental paradigm. Dorsal hFBOs were subjected to graded concussive injury at 25 DIV, and cell death was assessed at 1 and 3 days post-injury (dpi). **(C)** Representative TUNEL staining of intact and injured dorsal hFBOs exposed to mild, moderate, and severe intensities at 1 and 3 dpi. TUNEL-positive cells are shown in green and nuclei (DAPI) in blue. **(D, E)** Quantification of TUNEL-positive cells at 1 dpi **(D)** and 3 dpi **(E)** demonstrated an injury severity-dependent increase in cell death. TUNEL-positive cells were quantified as the percentage of TUNEL-positive cells relative to total DAPI-positive nuclei. Data are presented as mean ± SEM (n = 10 bins from 5 hFBOs per group, derived from 3 independent batches of 1 hiPSC line) and were analyzed by one-way ANOVA followed by Tukey’s multiple-comparisons post hoc test. ns, not significant; * p < 0.05, ** p < 0.01, *** p < 0.001, **** p < 0.0001.

We next examined injury-induced cell death at 80 DIV, when neuronal subtypes and dorsoventral identities are more established. For these experiments, dorsal and ventral hFBOs were subjected to moderate injury, and cell death was assessed at 1 dpi (Supplementary Fig. 3A). Immunostaining revealed a marked increase in TUNEL-positive cells following injury in both dorsal and ventral hFBOs compared with intact controls (Supplementary Fig. 3B). Quantification showed a significant increase in TUNEL-positive cells following injury in both dorsal and ventral hFBOs (Dorsal: Intact, 7.49 ± 1.07%; Injured, 62.26 ± 1.39%; Ventral: Intact, 8.49 ± 0.72%; Injured, 58.97 ± 2.34%; mean ± SEM; n = 5 bins for intact and 8 bins for injured groups from 3 hFBOs/group derived from 3 independent batch from one hiPSC line; Dorsal Intact vs. Injured, p < 0.0001; Ventral Intact vs. Injured, p < 0.0001). However, regional identity had no significant effect, and there was no interaction between injury and regional identity (Data are presented as mean ± SEM; two-way ANOVA with Tukey’s post hoc test). Consistent with this, TUNEL positivity did not differ between dorsal and ventral hFBOs under either intact or injured conditions. Collectively, these findings demonstrate that concussive injury induces severity-dependent cell death in hFBOs, with higher injury intensities associated with greater TUNEL positivity at 25 DIV. Furthermore, substantial injury-induced cell death was observed at 80 DIV, indicating that more differentiated hFBOs remain susceptible to mechanical injury. Notably, injury responses were comparable between dorsal and ventral hFBOs at 80 DIV, suggesting that regional identity does not significantly influence injury-induced cell death at this developmental stage.

### Concussive Injury Enhances Neurogenic and Neuroimmune Signaling in Brain Organoids

To examine cellular responses to concussive injury, we analyzed dorsal and ventral hFBOs at 25 DIV, a developmental stage enriched in neural progenitor cells and active neurogenic programs. This stage was selected because it represents a proliferative, progenitor-dominant window prior to extensive neuronal subtype specification and network maturation, allowing direct assessment of injury-induced effects on stem cell proliferation without confounding influences from mature synaptic circuitry (Townshend et al. 2020; Elkabetz et al. 2008). Dorsal and ventral hFBOs at 25 DIV were subjected to moderate mechanical injury (2-2.5 psi) and compared with intact controls at 3 dpi.

To determine whether moderate concussive injury alters neural progenitor populations, we first examined SOX2 and Nestin expression in 25 DIV dorsal and ventral hFBOs at 3 dpi. Immunostaining revealed increased SOX2 and Nestin expression in injured organoids compared with intact controls (Fig. 6A, B), accompanied by higher percentages of SOX2-positive cells (Intact, 45.00 ± 1.84%; Injured, 58.20 ± 2.83%; p = 0.0029; n = 6 and 5 bins from 5 hFBO, derived from 3 independent batches of 1 hiPSC line, respectively) and Nestin-positive cells (Intact, 47.57 ± 2.55%; Injured, 62.11 ± 2.07%; p = 0.0020; n = 7 and 5 bins from 3 hFBOs, derived from 3 independent batches from 1 hiPSC line, respectively) relative to the total number of DAPI-positive nuclei (Fig. 6B; unpaired two-tailed t-tests). To investigate whether these changes were accompanied by increased cell proliferation, we next examined Ki67 expression. Injured hFBOs exhibited greater Ki67 immunoreactivity and a higher percentage of Ki67-positive cells compared with intact controls (Intact, 39.27 ± 1.34%; Injured, 55.37 ± 2.72%; p = 0.0003; n = 6 and 5 bins from 3 independent batches from 1 hiPSC line, respectively; unpaired two-tailed t-test; Fig. 6A, B). To explore a potential signaling mechanism associated with this proliferative response, we examined β-catenin, a central mediator of canonical Wnt signaling (Zhang et al. 2013; Woodhead et al. 2006). Immunostaining revealed greater β-catenin expression following injury, supported by an elevated percentage of β-catenin-positive cells relative to intact controls (Intact, 32.50 ± 3.21%; Injured, 59.01 ± 4.57%; p = 0.0006; n = 7 and 5 bins from 3 hFBOs, from 3 independent batches from 1 hiPSC line, respectively; unpaired two-tailed t-test; Fig. 6A, B). Together, these findings suggest that moderate concussive injury (2-2.5 psi) increases the proportion of cells expressing neural progenitor- and proliferation-associated markers in developing hFBOs. The association with elevated β-catenin expression suggests a role for Wnt signaling in the injury-associated proliferative response.

**Figure 6.**
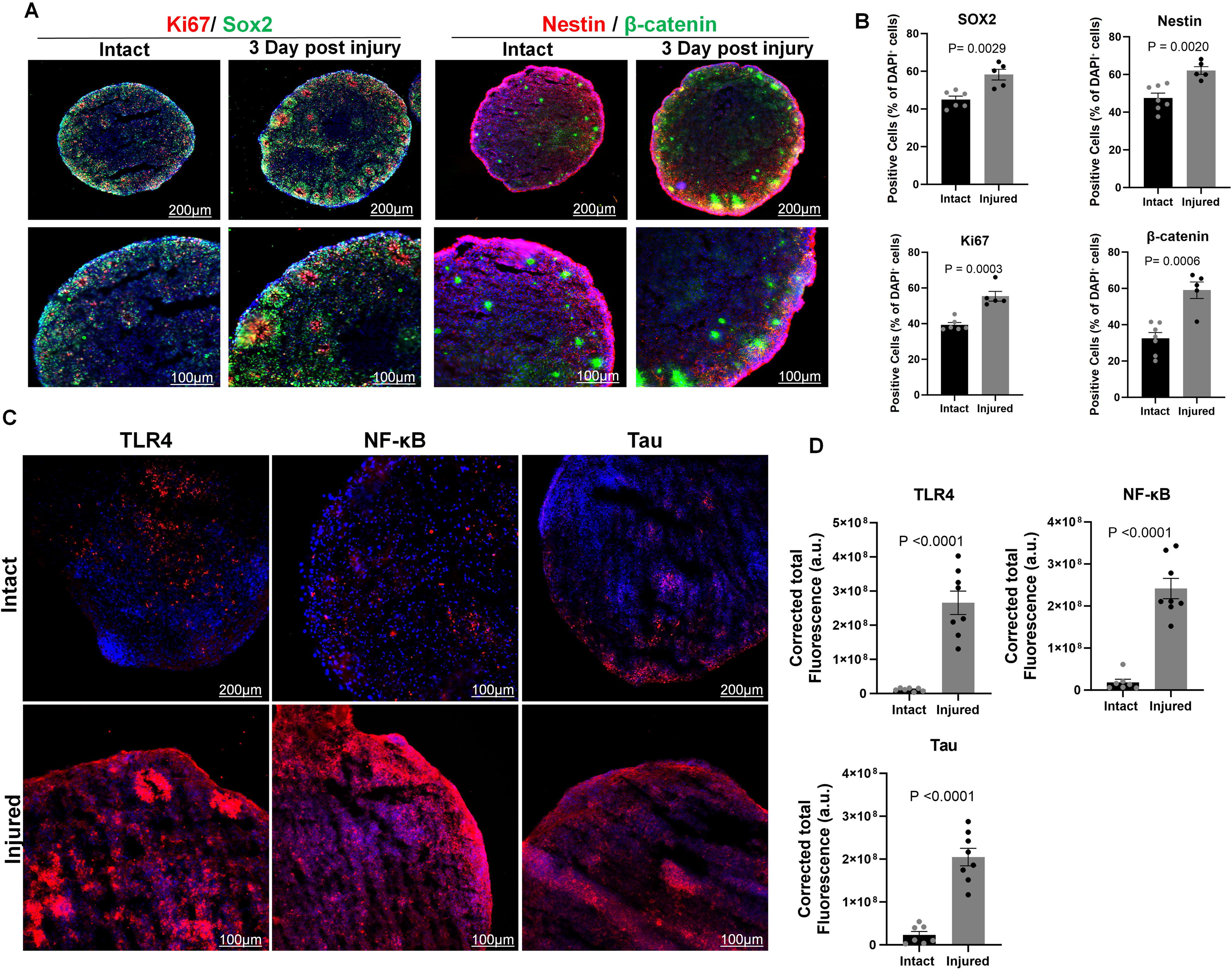
Concussive injury alters proliferative and immune signaling in human forebrain organoids. **(A)** Representative immunofluorescence images of dorsal hFBOs at 25 DIV comparing intact and injured organoids at 3 days post-injury (dpi; 2-2.5 PSI; moderate), showing SOX2 and Nestin expression, Ki67-positive proliferating cells, and β-catenin expression. Nuclei are counterstained with DAPI (blue). **(B)** Quantification shows increased SOX2-and Nestin-positive cells, Ki67-positive cells, and β-catenin-positive cells after injury. **(C)** Representative immunofluorescence images of TLR4, NF-κB, and Tau in intact and injured hFBOs at 80DIV examined 3 dpi. Nuclei are counterstained with DAPI (blue). **(D)** Quantification of fluorescence intensity demonstrates an increase in TLR4, NF-κB, and Tau expression following injury. Data are presented as mean ± SEM; for all groups, each dot represents an individual bin, with data obtained from 5 hFBOs per group across 3 independent batches derived from 1 hiPSC line and analyzed using unpaired, two-tailed t-tests. Fluorescence intensity is reported as corrected total cell fluorescence (CTCF; a.u.). Scale bars are indicated in each image.

We next investigated activation of injury-associated signaling pathwaysat 80 DIV and 3 dpi. To determine whether innate immune receptor signaling pathways implicated in rodent concussive injury are similarly altered in a human system, we first examined TLR4, an innate immune receptor previously shown to be upregulated after concussive brain injury in rodent models (Korgaonkar et al., 2020; Subramanian et al., 2026). Immunostaining revealed greater fluorescence intensity for TLR4 (Intact, 1.14 × 10⁷ ± 1.82 × 10⁶; Injured, 2.66 × 10⁸ ± 3.42 × 10⁷; n = 7 and 8 bins from 3 hFBO, derived from 3 independent batches from 1 hiPSC line, respectively; Fig. 6C, D), identifying increased TLR4 expression in injured hFBOs compared with intact controls. Because TLR4 can activate NF-κB signaling, we next examined NF-κB to test whether injury altered signaling pathways downstream of TLR4. Injured organoids showed greater NF-κB immunoreactivity (Fluorescence intensity in a.u, Intact: 1.85 × 10⁷ ± 7.46 × 10⁶; Injured: 2.42 × 10⁸ ± 2.43 × 10⁷; n = 7 and 8 bins from 3 independent hFBOs, respectively; Fig. 6C, D). To further test whether these injury-associated molecular changes were accompanied by alterations in neuronal integrity, we assessed Tau, a microtubule-associated protein that maintains neuronal cytoskeletal stability and is known to undergo pathological changes after traumatic brain injury. Immunostaining revealed greater Tau immunoreactivity after injury (Fluorescence intensity in a.u, Intact: 2.33 × 10⁷ ± 8.05 × 10⁶; Injured: 2.05 × 10⁸ ± 2.05 × 10⁷; n = 7 and 8 bins from 3 independent hFBOs, respectively; Fig. 6C, D). Together, these findings suggest that moderate concussive injury (2-2.5 psi) is associated with increased TLR4 and NF-κB expression, accompanied by elevated Tau immunoreactivity in developing hFBOs. Combined with the observed increase in progenitor- and proliferation-associated markers, these results indicate that moderate injury elicits concurrent proliferative and immune receptor-associated responses, alongside alterations in neuronal cytoskeletal protein expression.

### Calcium imaging reveals injury severity–dependent increases in calcium activity

To evaluate the functional consequences of mechanical injury on neuronal network activity, we performed live calcium imaging in hFBOs injured at 150 DIV. We selected this time point because our electrophysiological characterization showed that organoids at 150 DIV exhibit more mature intrinsic physiology and spontaneous synaptic activity, providing an appropriate stage for assessing network-level responses to injury. We focused on dorsal hFBOs because they showed more robust spontaneous excitatory synaptic activity at this stage, providing a more reliable platform to detect injury-evoked changes in activity dynamics. Calcium imaging was conducted 1 hour after mild, moderate, or severe injury to capture acute, early network responses before secondary cell death and tissue deterioration could confound functional readouts. Representative calcium heatmaps revealed a severity-dependent increase in fluorescence intensity, with the strongest signal after severe injury (Fig. 7A). To determine whether this increase reflected transient calcium activity or sustained calcium accumulation, we next examined calcium traces from individual cells. Intact hFBOs exhibited discrete calcium transients with recovery toward baseline, whereas mildly injured organoids showed broader and more prolonged calcium elevations in some cells. In contrast, moderate and severe injury produced predominantly sustained, non-recovering elevations or attenuation of calcium signals, depending on the region analyzed (Fig. 7B). These distinct temporal responses suggest that increasing injury severity shifts intracellular calcium dynamics from transient fluctuations toward sustained calcium accumulation, consistent with progressive intracellular calcium overload. To further characterize this response at the organoid level, we quantified global calcium fluorescence over a 150-second recording period using a whole-organoid region of interest (Fig. 7C). Intact and mildly injured hFBOs exhibited relatively stable fluorescence profiles, whereas moderate and severe injury produced sustained elevations in calcium signals (Fig. 7D). Quantification of peak fluorescence intensity revealed a significant effect of injury severity, with progressively higher intensities across groups (Intact, 44.38 ± 5.21; Mild, 59.37 ± 7.96; Moderate, 88.93 ± 5.53; Severe, 104.70 ± 12.17; mean ± SEM; n = 3 dorsal hFBOs/group; one-way ANOVA followed by Tukey’s post hoc test) (Fig. 7E). Compared with intact controls, peak intensity was significantly higher following moderate and severe injury, whereas mild injury produced a modest, non-significant increase. Together, these findings indicate that increasing injury severity is associated with progressive intracellular calcium accumulation, consistent with sustained calcium overload following moderate and severe concussive injury.

**Figure 7.**
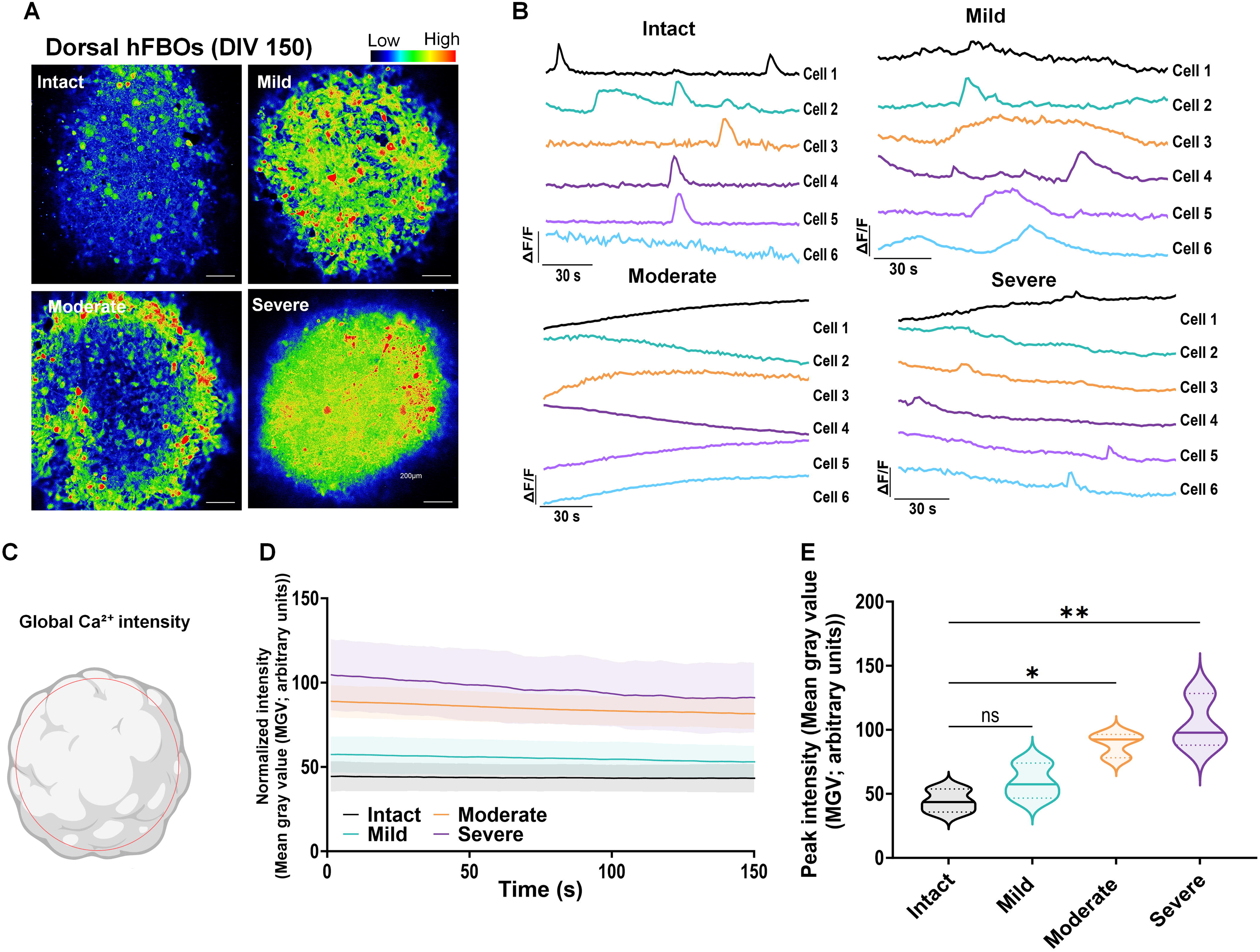
Injury severity-dependent calcium overload in dorsal human forebrain organoids. **(A)** Representative pseudocolored calcium fluorescence images of dorsal hFBOs at 150 DIV under intact conditions and following mild, moderate, or severe concussive injury, showing a progressive increase in calcium-dependent fluorescence with increasing injury severity. **(B)** Representative single-cell ΔF/F₀ traces from intact, mild, moderate, and severe injury groups. Intact and mildly injured hFBOs exhibit discrete calcium transients, whereas moderate and severe injury produce sustained, non-recovering elevations in intracellular calcium. **(C)** Schematic illustrating the whole-organoid region of interest (ROI) used to quantify global calcium fluorescence. **(D)** Global calcium fluorescence measured as mean gray value (MGV) over the 150-s recording period, demonstrating progressively elevated calcium signals with increasing injury severity. **(E)** Quantification of peak calcium fluorescence intensity (MGV) across intact, mild, moderate, and severe injury groups. Data are presented as mean ± SEM and were analyzed by one-way ANOVA followed by Tukey’s multiple-comparisons post hoc test (F(3,8) = 11.17, p = 0.0031). ns, not significant; * p < 0.05; **p < 0.01.

## Discussion

TBI is a complex, heterogeneous condition that triggers a cascade of cellular and molecular changes, contributing to both acute tissue damage and long-term neurological dysfunction (Brett et al. 2023). Although animal models have been instrumental in advancing our understanding of TBI pathophysiology, species-specific differences in brain development and neuronal maturation may limit the translation of preclinical findings to humans (Ciceri et al. 2024; Balakin et al. 2024). These limitations highlight the need for complementary human tissue-based experimental models that enable investigation of injury-induced molecular and cellular responses within the context of human neural development (Eichmüller and Knoblich 2022). Here, we established a regionally specified human iPSC-derived hFBO model with progressive cellular and electrophysiological maturation and adapted a controlled mechanical injury system to investigate the molecular and cellular consequences of concussive injury. Our findings demonstrate that dorsal and ventral transcriptional identities are detectable as early as 25 DIV, preceding the emergence of prominent region-specific neuronal populations at later developmental stages. By 80 DIV, regional cellular specification became evident, whereas intrinsic electrophysiological properties continued to mature through 150 DIV. Although intrinsic electrophysiological maturation was broadly comparable between dorsal and ventral hFBOs, we observed regional differences in excitatory synaptic activity. Using this platform, we demonstrate that concussive injury induces severity-dependent cell death, with developmental stage, rather than regional identity, influencing vulnerability to injury. At early developmental stages, injury increased neural progenitor proliferation and elevated TLR4 and NF-κB immunoreactivity, suggesting altered progenitor dynamics and injury-responsive signaling. In more mature hFBOs with functional neuronal circuits, injury induced severity-dependent calcium dysregulation, characterized by sustained calcium elevations following higher-severity injury. Together, these findings establish a developmentally and regionally characterized human forebrain organoid platform for investigating how developmental stage and regional identity influence responses to concussive injury.

The developing forebrain is organized into two major domains, the dorsal pallium and ventral subpallium, which give rise to neuronal populations enriched in glutamatergic excitatory and GABAergic inhibitory neurons, respectively (Marín and Müller 2014). We find that hFBOs show regional transcriptomic differentiation by 25 DIV, with cellular differentiation established by 80 DIV, consistent with the temporal and regional developmental programs of hFBOs at transcriptional and cellular levels. Expression of NKX2-1, required for the development of major GABAergic neuronal subtypes, occurrs by day 25, followed by the emergence of GABAergic neurons expressing GABA and GAD67 by day 60 and interneuron subtype markers, including SST and PV, by day 109 (Birey et al. 2017). Similarly, regionally patterned cerebral organoids develop distinct dorsal and ventral identities, with dorsal populations enriched for cortical glutamatergic neuronal markers and ventral populations enriched for progenitor and inhibitory neuronal lineage markers including SST- and PV-expressing interneuron subtypes. Consistent with these findings, our hFBOs exhibited a similar regional pattern, with dorsal cortical progenitor identity and ventral inhibitory neuronal differentiation (Bagley et al. 2017). Transcriptomic and epigenomic analyses further revealed distinct molecular programs associated with cortical and medial ganglionic eminence organoid development (Xiang et al. 2017), while single-cell transcriptomic studies demonstrated progressive neuronal differentiation and diversification of region-specific cellular populations over time (Kanton et al. 2019; Velasco et al. 2019). More recent investigations of FGF8-mediated regional patterning have further distinguished pallial populations enriched in glutamatergic neuronal markers, including EMX1, TBR1, and SLC17A7, from subpallial populations enriched in GABAergic lineage markers, including ASCL1, DLX1/2, and GAD1/2 (Bertacchi et al. 2024). Consistent with these findings, our transcriptional profiling revealed distinct dorsal and ventral identities as early as 25 DIV, when both organoid populations remained predominantly composed of neural progenitors. These early transcriptional differences preceded the emergence of prominent region-specific neuronal populations at 80 DIV, with dorsal hFBOs enriched in excitatory cortical neuronal markers, including CTIP2, and ventral hFBOs enriched in inhibitory neuronal markers, including GABA and SST. Interestingly, we detected robust PV immunoreactivity in ventral hFBOs at 150 DIV, indicating the emergence of an additional inhibitory neuronal subtype during prolonged culture. Although generating of PV-expressing interneurons has been challenging in human brain organoid models, recent studies have reported PV expression in MGE-derived organoids by four months, accompanied by the expression of Kv3.1, a voltage-gated potassium channel associated with the rapid repolarization and high-frequency firing characteristic of mature PV interneurons (Varela et al. 2025). Our detection of PV-positive cells at 150 DIV further supports the capacity of ventral hFBOs to generate diverse inhibitory neuronal populations during extended differentiation. Together, these findings suggest that pallial and subpallial molecular identities are established during early progenitor-dominant stages and become increasingly evident through the differentiation of glutamatergic and GABAergic neuronal populations, respectively. Importantly, defining these developmental transitions within the same experimental platform provides a framework for investigating how regional identity and developmental stage contribute to cellular responses following concussive injury.

Functional maturation is a critical feature of human brain organoid development, reflecting the progressive acquisition of neuronal excitability and synaptic connectivity. Previous studies have demonstrated the development of action potential firing and functional synaptic transmission in human cortical organoids over 130 DIV (Pasca et al. 2015), and prolonged organoid culture has been associated with increased synaptic organization and neuronal network activity (Quadrato et al. 2017). Studies using multielectrode arrays have demonstrated progressive emergence of synchronized network activity and increasingly complex oscillatory patterns during organoid maturation (Trujillo et al. 2019). Although these studies established the capacity of human brain organoids to develop functional neuronal networks, population-level recordings do not directly resolve the intrinsic membrane properties and synaptic inputs of individual neurons. In this context, our patch-clamp characterization provides a complementary perspective by comparing intrinsic electrophysiological maturation and excitatory synaptic transmission in regionally specified dorsal and ventral hFBOs across defined developmental stages. The developmental changes in resting membrane potential, input resistance, and action potential properties observed between 80 and 150 DIV are consistent with progressive maturation of neuronal membrane conductance, potentially involving changes in voltage-gated sodium and potassium channel function and membrane permeability (Wu et al. 2020). Importantly, the emergence of spontaneous excitatory postsynaptic currents at 150 DIV, following the exclusive occurrence of evoked synaptic responses at 80 DIV, suggests that functional synaptic connections are established before the development of spontaneous excitatory transmission. Notably, dorsal and ventral neurons exhibited broadly comparable intrinsic electrophysiological maturation, whereas dorsal hFBOs displayed more frequent spontaneous excitatory synaptic events at 150 DIV. This distinction suggests that intrinsic neuronal maturation and synaptic functional development are not necessarily coupled to the same extent across regional identities. Despite comparable intrinsic electrophysiological maturation, dorsal hFBOs exhibited greater excitatory synaptic activity than ventral hFBOs, potentially reflecting their enrichment in glutamatergic neurons. These findings suggest that regional identity influences synaptic function and provide a foundation for investigating neuronal responses to concussive injury.

Previous studies have demonstrated the potential of human brain organoids to model traumatic brain injury using different mechanical approaches, including controlled cortical impact, compression, and ultrasound-induced injury (Ramirez et al. 2021; Lai et al. 2024). These models have provided important insights into injury-induced cellular damage and molecular responses in human neural tissue. Mild and moderate mechanical loading of 4-week-old human cerebral organoids has been shown to alter gene expression and calcium signaling, highlighting the utility of organoid models for examining responses to mechanical stress (Beltrán et al. 2023). Building on these advances, we adapted a controlled mechanical injury system to induce concussive injury in regionally specified human forebrain organoids at distinct developmental stages. By incorporating dorsal and ventral regional identities and progressive cellular and functional maturation, our model provides an opportunity to investigate injury responses in different developmental and cellular contexts. Our findings demonstrate that increasing injury severity led to progressively greater cell death in early-stage hFBOs, highlighting the sensitivity of developing human neural tissue to the magnitude of mechanical injury. Similar associations between injury severity and the extent of neuropathology have been documented in human clinical and postmortem studies (Mckee and Daneshvar 2015). Interestingly, the initial increase in TUNEL positivity following lower-pressure injury observed 1 dpi was no longer evident at 3 dpi, whereas higher injury pressures were associated with persistently elevated TUNEL positivity. This pattern suggests that mild injury may produce a more limited cellular damage response, consistent with observations in rodent models of TBI (Toth et al. 1997; Raghupathi et al. 2002). In contrast, the sustained elevation of TUNEL positivity following higher injury pressures may reflect more extensive or prolonged cellular damage, potentially involving secondary injury processes that extend beyond the initial mechanical insult. Furthermore, the comparable injury-induced cell death observed in dorsal and ventral hFBOs at 80 DIV suggests that differences in regional neuronal specification do not necessarily translate into distinct susceptibility to mechanical injury under the conditions examined. Although differential vulnerability among specific neuronal populations cannot be excluded, these findings suggest that overall injury-associated cell death may be similar across dorsal and ventral forebrain identities. Together, these observations support the utility of regionally specified hFBOs as a human-tissue platform for investigating injury severity-dependent cellular damage and the temporal evolution of injury responses across distinct developmental and regional contexts.

Traumatic brain injury has been shown to influence neural progenitor proliferation and neurogenic responses in rodent models (Dash, Mach, and Moore 2001; Kernie, Erwin, and Parada 2001; Neuberger, Swietek, et al. 2017), with similar changes in progenitor-associated populations reported in human brain tissue following TBI (Zheng et al. 2013). Recent studies show that mechanical compression of young (30 DIV) human brain organoids transiently increases SOX2-positive neural precursor cell expression and reduces neuronal differentiation, but not when injury occurs after 50 DIV (Lampersperger et al. 2026). Their transcriptomic analyses of injured organoids 30 DIV revealed changes in WNT-associated developmental programs, suggesting that mechanical forces can influence signaling pathways involved in neural progenitor regulation and regional patterning. Our results extend these findings by demonstrating that a controlled concussive injury model in developing hFBOs at 25 DIV increased SOX2, Nestin, and Ki67 immunoreactivity and elevated β-catenin expression, a downstream effector of the WNT signaling pathway. Our findings indicate that concussive injury influences progenitor maintenance and proliferative activity during early neural development, potentially altering the balance between progenitor expansion and neuronal differentiation. The accompanying increase in β-catenin immunoreactivity is particularly noteworthy, as it raises the possibility that canonical Wnt/β-catenin signaling may contribute to injury-associated changes in progenitor dynamics, consistent with its established role in regulating neural progenitor proliferation and differentiation (Munji et al. 2011). Although these responses may represent an adaptive attempt to maintain progenitor populations after injury, persistent alterations in progenitor regulation could also disrupt normal neuronal differentiation and maturation. Importantly, increased progenitor and proliferation-associated marker expression does not necessarily indicate successful neurogenesis or neuronal replacement, and elevated β-catenin immunoreactivity alone does not establish canonical Wnt pathway activation. Future studies examining Wnt signaling activity, progenitor cell-cycle dynamics, and subsequent neuronal differentiation will be necessary to determine whether these injury-associated responses support neural repair or contribute to altered developmental trajectories.

We find that concussive injury in hFBO at 80 DIV increased TLR4 and NF-κB expression despite the absence of a conventional microglial population. Although TLR4 is typically associated with microglia-mediated innate immune responses, our previous studies have demonstrated neuronal TLR4 upregulation following concussive brain injury and identified its role in regulating neuronal excitability (Korgaonkar, Nguyen, et al. 2020). In experimental TBI, TLR4 expression and neural progenitor proliferation increase in parallel (Ye et al. 2014). However, whether TLR4 signaling causally drives the post-injury increase in NSCs remains to be determined. Increases in neuronal TLR4 have been shown to be central to synaptic and circuit reorganization by regulating MMP-9, an extracellular matrix remodeling enzyme (Subramanian et al. 2026). These observations suggest that hFBOs, like murine models (Korgaonkar, Nguyen, et al. 2020; Subramanian et al. 2026) respond to injury by activating TLR4 signaling beyond conventional microglial-mediated pathways. Although previous human brain organoid injury studies have identified alterations in immune-associated transcriptional programs (Beltrán et al. 2023), the contribution of TLR4-associated signaling to mechanical injury responses in human neural tissue remains unexplored. In this context, our observation of increased TLR4 and NF-κB immunoreactivity following concussive injury in hFBOs lacking microglia suggests that injury to hFBOs neural cells may exhibit injury-associated changes in innate immune receptor signaling. Although NF-κB is downstream of TLR4 signaling, multiple pathways can enhance NF-κB. Future investigations of cell type-specific TLR4 expression, NF-κB activation, and downstream signaling are needed to determine whether these responses contribute to injury-induced cellular damage or represent adaptive signaling mechanisms in developing human neural tissue.

The increase in Tau immunoreactivity is also relevant given the association between TBI and Tau-related pathology. A prior study reported changes in the phosphorylated-to-total Tau ratio following mechanical injury in human brain organoids, demonstrating the potential of organoid models to capture injury-associated alterations in Tau biology (Lai et al. 2024). Our findings extend these observations by demonstrating increased Tau immunoreactivity in regionally specified hFBOs following concussive injury. Together, the injury-associated increases in TLR4, NF-κB, and Tau highlight molecular responses to mechanical injury in developing human neural tissue and provide a foundation for future studies investigating their potential contributions to long-term alterations in neuronal development and function.

Beyond injury-associated molecular changes, disruption of intracellular calcium homeostasis represents an important mechanism of secondary neuronal dysfunction following TBI. Calcium is essential for neuronal excitability, synaptic transmission, and intracellular signaling, whereas excessive or sustained calcium elevations can contribute to mitochondrial dysfunction, activation of calcium-dependent proteases, and neuronal injury (Brini et al. 2014; Weber 2012). Previous studies have shown that mechanical injury alters calcium levels in young (30 DIV) human cerebral organoids (Beltrán et al. 2023), indicating that organoid models can reproduce injury-severity-dependent calcium responses. Building on these observations, we examined injury-induced calcium dynamics in more functionally differentiated hFBOs at 150 DIV, a stage at which whole-cell patch-clamp recordings independently demonstrated spontaneous neuronal activity and synaptic transmission. The sustained calcium elevations we observed following moderate and severe injury suggest that mechanical stress can disrupt calcium homeostasis in human neural tissue with established electrophysiological activity. Unlike the discrete calcium transients observed in intact and mildly injured organoids, prolonged elevations following higher injury severity may reflect impaired cytotoxic calcium increases, which could contribute to secondary cell death at stronger injuries. Together, these findings extend previous organoid-based injury studies by linking severity-dependent calcium dysregulation to a developmentally characterized human forebrain model with demonstrated spontaneous neuronal and synaptic activity, providing an opportunity to investigate how mechanical injury affects the physiology of functionally differentiated human neural tissue.

A major strength of this study is the integration of regionally specified human forebrain organoids, progressive functional maturation, and controlled mechanical injury, enabling investigation of injury responses across distinct developmental stages in a human neural context. However, several limitations warrant consideration. Although hFBOs reproduce key aspects of human forebrain development, they lack vascularization, resident microglia, and the full cellular and structural complexity of the intact brain, limiting their ability to capture systemic inflammatory and vascular responses to TBI. Furthermore, while injury-associated changes in progenitor and proliferation markers were observed, the long-term consequences for neuronal differentiation and maturation remain to be determined. Similarly, calcium imaging identified sustained elevations following moderate and severe injury, but the underlying mechanisms and their consequences for neuronal and synaptic function require further investigation. Despite these limitations, this platform provides a complementary approach for examining cellular and functional responses to concussive injury in developing human neural tissue.

Collectively, this study establishes a human forebrain organoid platform for investigating the cellular and functional consequences of concussive injury across developmental stages. We demonstrate severity-dependent cellular damage, injury-associated changes in neural progenitor and signaling responses, and sustained calcium elevations following moderate and severe injury in mature organoids. By combining regional specification, progressive functional maturation, and controlled mechanical injury, this accessible platform complements existing TBI models and provides opportunities to investigate how developmental stage and neuronal identity influence injury responses. Future studies incorporating longitudinal functional assessments and dorsal– ventral assembloids may further elucidate the consequences of concussive injury for neuronal maturation, circuit development, and regional vulnerability.

## Acknowledgements

This work was supported by NIH/NINDS grants R01NS069861 and R01NS097750 to V.S., the CIRM TRANSCEND Fellowship (EDUC4-12752) to R.J., NIH/NIDCR grants R01DE025330 and K18DE032513 to N.z.N, and NIEHS T32 traineeship to P.O.B. (T32ES018827).

## Author contributions

R.J. performed experiments, including organoid generation and culture, injury induction, electrophysiological recordings, immunohistochemistry, and TUNEL assays; collected and analyzed data; prepared figures; and drafted the manuscript. P.D.O. performed calcium imaging experiments and analysis. C.T.J. generated graphical illustrations. E.C. assisted with cell culture. R.C. performed blind data analysis. P.O.B. contributed to data visualization. I.N. reviewed and edited the manuscript. N.I.Z.N. provided supervision and reviewed and edited the manuscript. V.S. conceived and designed the study, supervised and managed the project, acquired funding, contributed to data interpretation, and revised and finalized the manuscript. All authors reviewed and approved the final manuscript.

## Declaration of interests

The authors declare no competing interests.

## Notes

### Competing Interest Statement

The authors have declared no competing interest.

