## Supplementary Figures and Tables for "Concussive Injury in Human Forebrain Organoids Alters Neurogenesis and Neuroimmune Signaling"

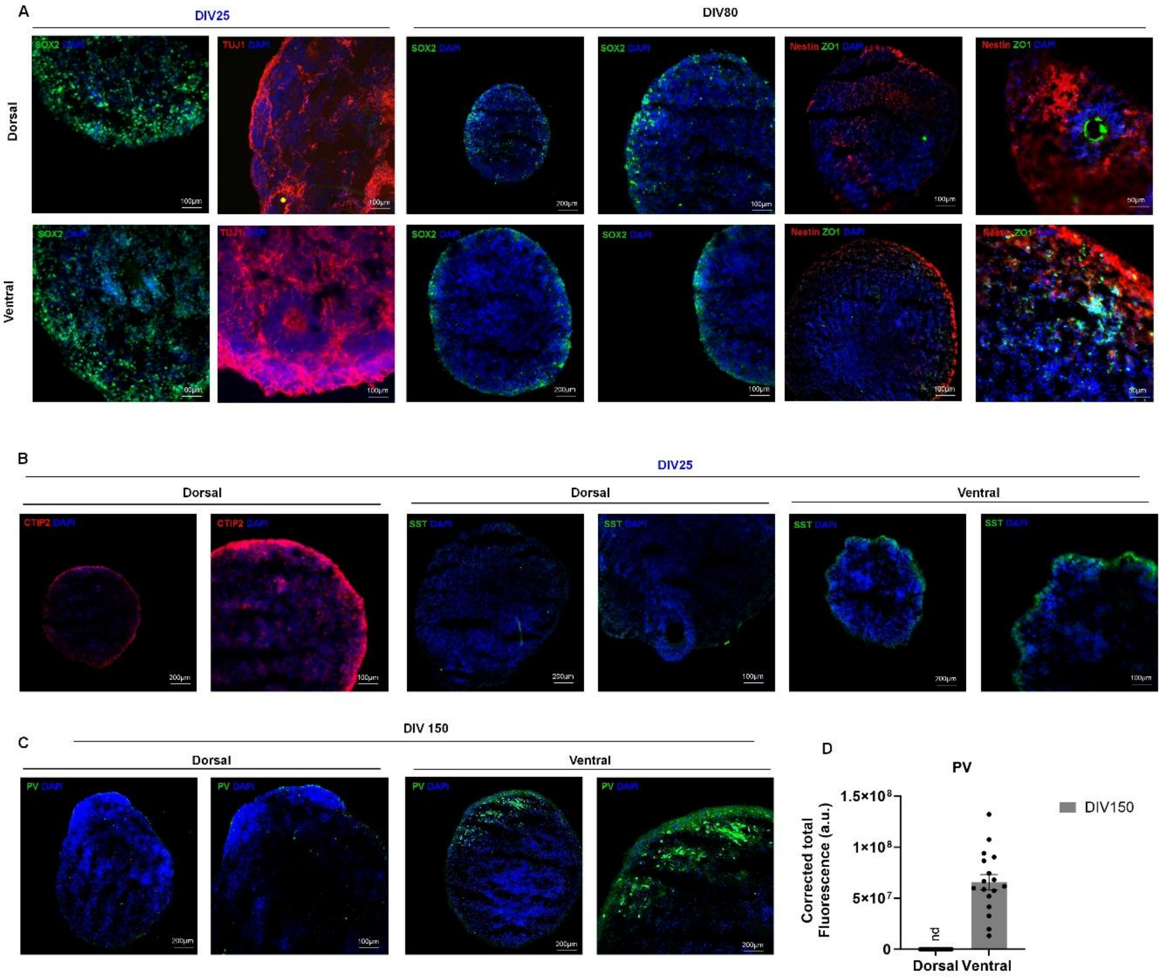


**Supplementary Figure 1. Temporal and regional characterization of marker expressions in hFBOs. (A)** Representative images show SOX2 and TUJ1 expression in dorsal and ventral hFBOs at 25 and SOX2 expression at 80 DIV. Representative images show Nestin and ZO-1 expression in sections from dorsal and ventral hFBOs at 80 DIV, but expression was reduced compared with early stages (see Fig. 1 for expression at 25 DIV). **(B)** Representative images of CTIP2 and SST expression at 25 DIV. SST expression is shown in dorsal and ventral hFBO. CTIP2 was not detected in ventral hFBOs. **(C)** Representative images show expression of parvalbumin (PV) in dorsal and ventral hFBOs at 150 DIV. **(D)** Summary plot comparing PV fluorescence intensity between dorsal and ventral organoids at 150 DIV.

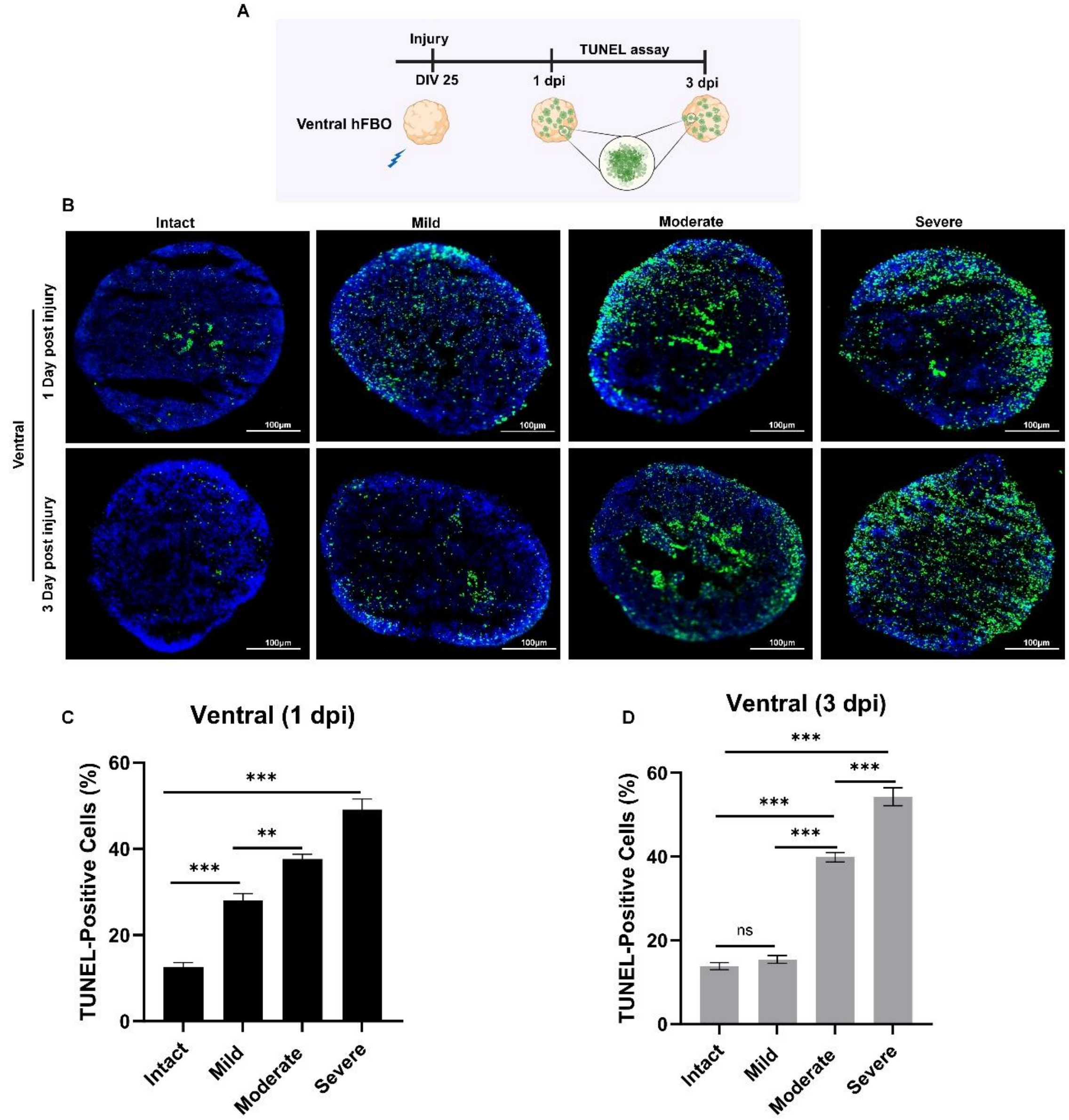


**Supplementary Figure 2. Injury severity-dependent cell death in ventral human forebrain organoids.** **(A)** Schematic of the experimental paradigm. Ventral hFBOs were subjected to graded concussive injury at 25 DIV, and cell death was assessed at 1 and 3 days post-injury (dpi). **(B)** Representative TUNEL staining of intact and injured ventral hFBOs exposed to mild, moderate or severe injury at 1 and 3 dpi. TUNEL-positive cells are shown in green and nuclei (DAPI) in blue. **(C, D)** Quantification of TUNEL-positive cells at 1 dpi **(C)** and 3 dpi **(D)** demonstrates an injury severity-dependent increase in cell death. n = 8 bins from 5 hFBOs per group, derived from 3 independent batches of 1 hiPSC line. Data are presented as mean ±

SEM and were analyzed by one-way ANOVA followed by Tukey's multiple-comparisons post hoc test. ns, not significant; **$p < 0.01$; ***$p < 0.001$; ****$p < 0.0001$.

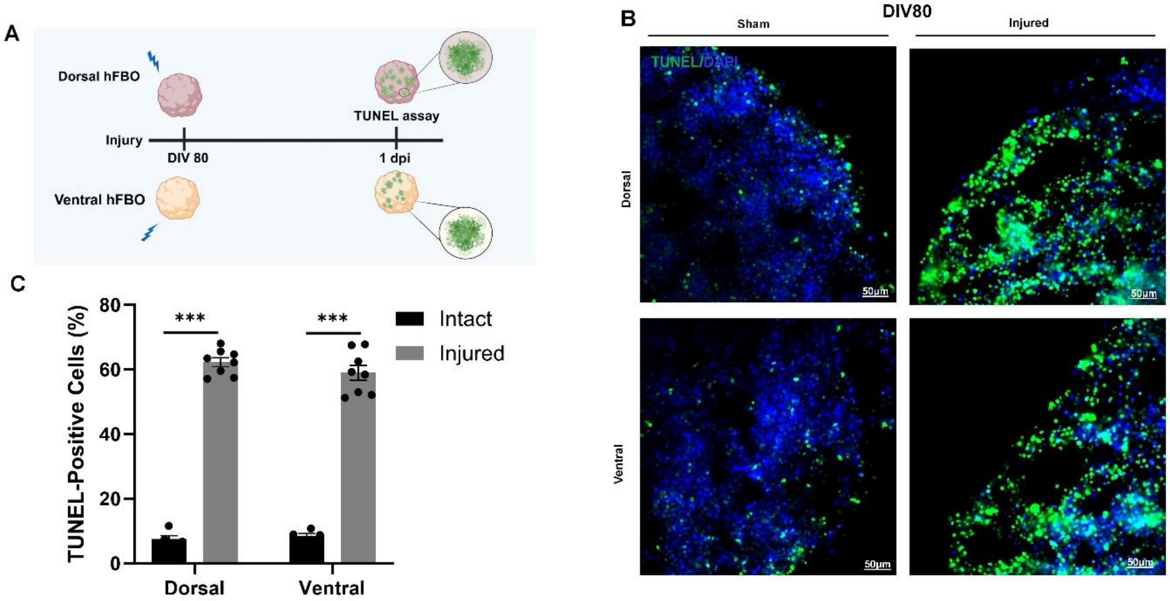


**Supplementary Figure 3. Concussive injury induces comparable cell death in dorsal and ventral human forebrain organoids at 80 DIV. (A)** Schematic of the experimental paradigm. Dorsal and ventral hFBOs at 80 DIV were subjected to concussive injury, and cell death was assessed by TUNEL staining at 1 day post-injury (dpi). **(B)** Representative TUNEL staining of intact and injured dorsal and ventral hFBOs. TUNEL-positive cells are shown in green and nuclei (DAPI) in blue. **(C)** Quantification of TUNEL-positive cells demonstrates a significant increase in cell death following injury in both dorsal and ventral hFBOs, with no significant difference between regional identities. TUNEL-positive cells were quantified as the percentage of TUNEL-positive cells relative to total DAPI-positive nuclei. Data are presented as mean ± SEM; n = 5 bins for intact and 8 bins for injured groups from 3 hFBOs/group derived from 3 independent batches across 1 hiPSC and were analyzed by two-way ANOVA with injury status and regional identity as factors, followed by Tukey's multiple-comparisons post hoc test. *** $P < 0.001$.

**Supplementary table 1. Antibodies**

| Antibodies | Source | Identifier |
|---|---|---|
| Mouse anti-SOX2 | Abcam | Cat# ab79351; RRID: AB_10710406 |
| Rabbit anti-Nestin | Sigma-Aldrich | Cat# SAB4200347 |
| Mouse anti-ZO-1 | Invitrogen | Cat# 33-9100; RRID: AB_2533147 |
| Rabbit anti-PAX6 | MBL International | Cat# PD022 |
| Mouse anti-S100β (clone 4C4.9) | Invitrogen | Cat# MA5-12969; RRID: AB_10983883 |
| Mouse anti-βIII-tubulin (TUJ1) | BioLegend | Cat# 801202; RRID: AB_2313773 |
| Rat anti-CTIP2 | Abcam | Cat# ab18465; RRID: AB_2064130 |
| Rabbit anti-GABA | Invitrogen | Cat# PA5-32241; RRID: AB_2549714 |
| Rabbit anti-Parvalbumin (PV) | Invitrogen | Cat# PA1-933; RRID: AB_2173898 |
| Rabbit anti-Somatostatin (SST) | Invitrogen | Cat# PA5-82678; RRID: AB_2789834 |
| Rabbit anti-Ki67 | Thermo Fisher Scientific | Cat# MA5-14520; RRID: AB_10979488 |
| Mouse anti-β-catenin (clone 9F2) | Invitrogen | Cat# MA1-2001; RRID: AB_326078 |
| Mouse anti-TLR4 (clone 25) | Santa Cruz Biotechnology | Cat# sc-293072; RRID: AB_10611320 |
| Mouse anti-NF-κB | Santa Cruz Biotechnology | Cat# sc-8008; RRID: AB_628017 |
| Mouse anti-Tau | EMD Millipore | Cat# 05-803; RRID: AB_310013 |
| Goat anti-mouse IgG Alexa Fluor 488 | Invitrogen | Cat# A-11001; RRID: AB_2534069 |
| Goat anti-mouse IgG Alexa Fluor 568 | Invitrogen | Cat# A-11031; RRID: |
| Goat anti-rabbit IgG Alexa Fluor 488 | Invitrogen | Cat# A-11008; RRID: AB_143165 |
| Goat anti-rabbit IgG Alexa Fluor 594 | Invitrogen | Cat# A-11037; RRID: AB_2534095 |
| Goat anti-rat IgG Alexa Fluor 488 | Invitrogen | Cat# A-11006; RRID: AB_2534074 |
